# Acute opioid responses are modulated by dynamic interactions of *Oprm1* and *Fgf12*

**DOI:** 10.1101/2022.03.11.483993

**Authors:** Paige M. Lemen, Yanning Zuo, Alexander S. Hatoum, Price E. Dickson, Guy Mittleman, Arpana Agrawal, Benjamin C. Reiner, Wade Berrettini, David G. Ashbrook, Mustafa Hakan Gunturkun, Xusheng Wang, Megan K. Mulligan, Caleb J. Browne, Eric J. Nestler, Francesca Telese, Robert W. Williams, Hao Chen

## Abstract

Exploring the molecular genetic cascades responsible for behavioral responses to opioids can improve our understanding of drug use initiation. We generated high-precision time-series data for 105 morphine-and naloxone-related traits across ∼700 young adult BXD mice (64 diverse strains and both sexes) for 3 hours after a single morphine injection. Variations in responses were mapped using high-precision genome sequencing-based genotypes. The initial locomotor responses to morphine mapped precisely to the µ opioid receptor gene (*Oprm1*) on chromosome (Chr) 10 with a peak linkage of 12.4 (-logP, genome-wide significance level is approximately 3.8). The *B* allele inherited from C57BL/6J was associated with up to 60% higher activity. This effect climaxed at 75 min but was exhausted by 160 min. A second major modulator of opioid-induced locomotion emerged after approximately 100 min. This locus, also associated with a high *B* allele, was located on Chr 16 with peak linkage of 10.6 (-logP) in females. This locus included only one compelling candidate, fibroblast growth factor 12 (*Fgf12*), a 600 Kb gene that controls sodium current kinetics at the axon hillock. A strong and transient epistatic interaction existed between the *Oprm1* and *Fgf12* loci during a short time window (45-75 min). The combination of a *B* haplotype at *Oprm1* with a *D* haplotype from DBA/2J at *Fgf12* was associated with unusually high activity. In a complementary study in heterogeneous stock rats, we demonstrated that *Oprm1* and *Fgf12* were co-expressed in one specific subtype of Drd1^+^ medium spiny neuron. A Bayesian network analysis supported an *Oprm1*-to-*Fgf12* network that involves a MAP kinase cascade that modulates *FGF12* phosphorylation and locomotor activation. *OPRM1* and *FGF12* networks in human GWAS data highlight enrichment of signals associated with substance use disorder. This study represents the first demonstration of a time-dependent epistatic interaction modulating drug response in mammals and the first linkage of *Fgf12* to opioid-induced behavior.

## Introduction

Understanding the molecular genetics of opioids is crucial for treatment of opioid use disorder (OUD). Great progress has been made recently in genome-wide association studies (GWAS) of OUD. Genetic variants of several genes have reached genome-wide significance, such as *KCNC1*, *KCNC2* (Gelernter et al., 2014), *CNIH3* (Nelson et al., 2016), *RGMA* (Cheng et al., 2018), *KDM4A* (Sanchez-Roige et al., 2021), and *OPRM1* (Zhou et al., 2020). Recently, Deak and colleagues (2022) have identified an additional set of 19 independent risk loci for OUD.

Most of these variants explain a relatively small fraction of the heritability of OUD and require validation.

The most replicated gene among these studies is the opioid receptor gene *OPRM1* (Deak et al., 2022; Gaddis et al., 2022; Zhou et al., 2020), which encodes the μ opioid receptor (MOR).

Morphine, the prototypic MOR agonist, inhibits adenylyl cyclase and decreases the transmission of nociceptive information (Yam et al., 2018). Activation of MOR also increases dopaminergic neuronal transmission to the nucleus accumbens (NAc) by inhibiting GABAergic interneurons in the ventral tegmental area (VTA) (De Vries and Shippenberg, 2002). Coexpression of both MORs and dopamine D1 receptors (DRD1)—but not D2 receptors (DRD2)— is required for the initial locomotor response to morphine (Severino et al., 2020). MOR activation in striatal neurons is also a sufficient signal of opioid reward (Cui et al., 2014). However, the downstream molecular networks associated with *OPRM1* activation and signaling remain incompletely understood.

The lack of large and well phenotyped populations is a major impediment to further discovery of specific genes and mechanisms that modulate different phases of OUD (Berrettini, 2017). Compared to human GWAS, genetic mapping using model organisms requires a much smaller sample size. Phenotypes can be measured more accurately and studied mechanistically with better control over environmental variables. Large families of recombinant inbred strains (RI) are especially powerful for identifying genetic drivers responsible for trait variation, such as responses to opioids. These families are made by crossing two or more inbred strains, followed by intercrossing and inbreeding progeny for more than 20 generations (Williams et al., 2001).

The BXD RI family of mice was created by crossing C57BL/6J (B6) with DBA/2J (D2) (Ashbrook et al., 2021; Taylor et al., 1999, 1973). This family has been used to study the effects of alcohol and many psychoactive drugs (Belknap et al., 1993; Crabbe, 1998; Crabbe et al., 1983; Miner and Marley, 1995), along with numerous other molecular and complex traits.

The original data were described in Philip et al. (2010), who identified the *Oprm1* locus. Here, we significantly extended these analyses by using whole-genome sequence-based genetic maps (Ashbrook et al., 2021) and the GEMMA linear mixed model (LMM) (Zhou and Stephens, 2012) for mapping, which improves mapping power and controls for population structure better than previously used methods. In addition to *Oprm1*, our analysis identified a new morphine response locus on chromosome 16 (Chr 16). We exploited complementary rodent data sets—both whole brain proteomics and single-nuclei RNAseq—and identified a new candidate gene—fibroblast growth factor 12 (*Fgf12*). We found a strong and transient epistatic interaction between the *Oprm1* and *Fgf12* loci in a short time window after morphine injection.

The snRNA-seq data highlights a single subtype of *Drd1*-positive cell type in which *Oprm1* and *Fgf12* are coexpressed. We then created a probabilistic causal model formalizing a molecular scheme in which gene variants modulate key signaling molecules to control locomotion after morphine injection. Lastly, we integrated rodent data with comparable human gene expression to provide a better translational context for understanding the molecular mechanisms and cellular cascades that may underlying initial variation in human OUD responses.

## Results

### QTL mapping of morphine-induced locomotor responses

A total of 64 fully inbred strains from the BXD family (Ashbrook et al., 2021) were used. Each strain was represented by an average of six males and six females (Philip et al., 2010).

Locomotion was recorded for 180 min after an acute morphine injection, with movement data binned for each 15 min period. All data were generated by co-authors (PED, GM). Data for the 10th time bin (135–150 min) were lost. Variation in locomotor responses revealed a skewed distribution (Supplementary Figure 1). We therefore quantile normalized data (Supplementary Figure 2) for most analyses, although we note that linkage statistics are robust with respect to this transformation.

QTL linkage was computed using a subset of ∼7,000 informative sequenced-based markers selected via linkage disequilibrium pruning and using the latest version of GEMMA v0.98.5 (Prins et al., github.com/genetics-statistics/GEMMA) as implemented in GeneNetwork.org. The genome-wide –logP significance threshold (*p* <0.05), determined by 1,000 permutations, is approximately 3.77 for the BXD family. A list of genome-wide significant loci for both morphine-induced locomotion and naloxone-induced withdrawal was summarized in Table 1, with more details provided in Supplementary Table 1. We analyzed both sexes separately and jointly to identify potential sex-specific genetic modifiers of morphine response, because significant sex by strain interaction was detected in our prior analysis (Philip et al., 2010).

**Table 1.**
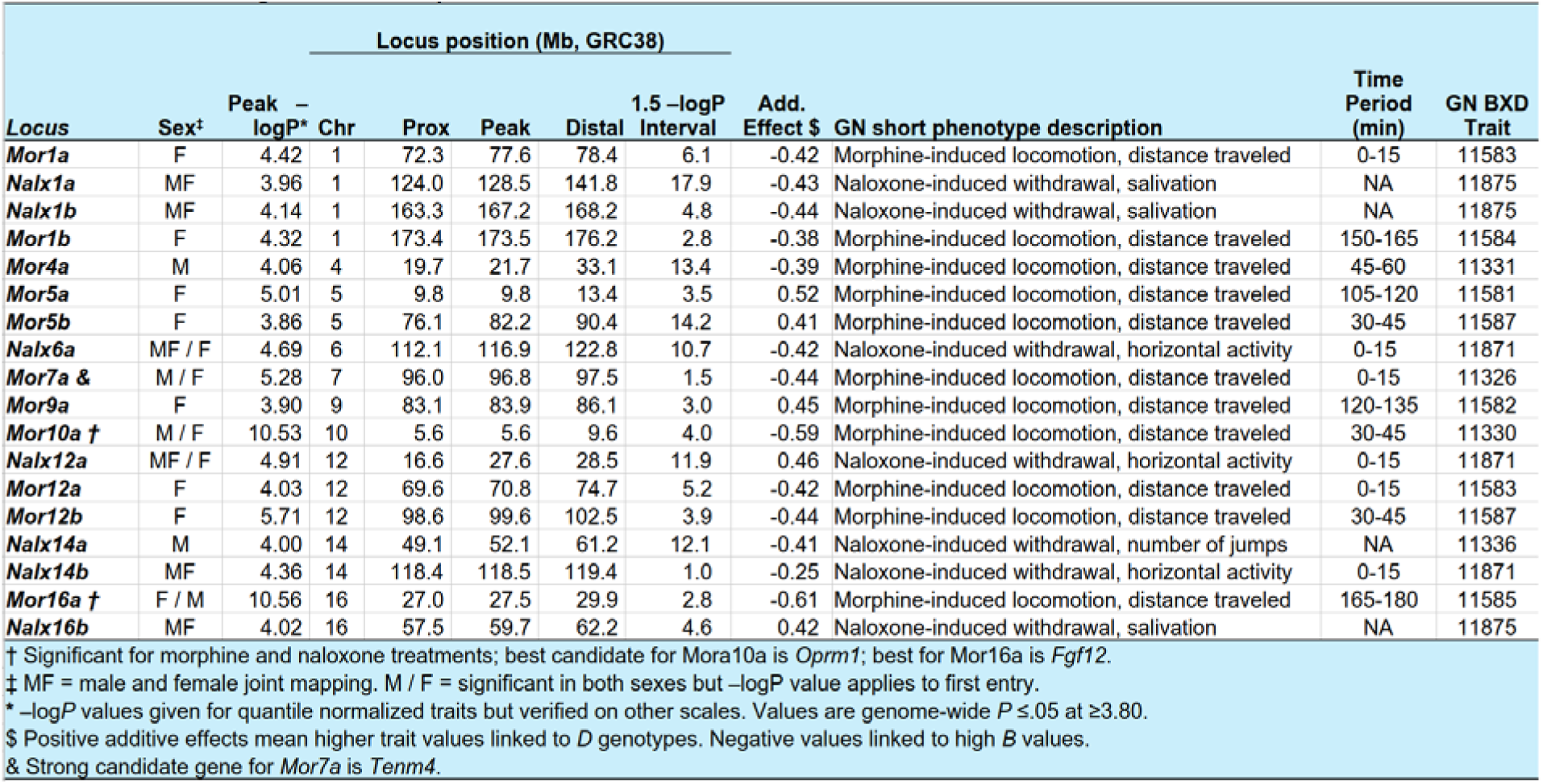
Genome-Wide Significant Loci for Morphine-Induced Locomotion or Naloxone-Induced Withdrawal.

Throughout the first two hours after injection of morphine (0–135 min bins) we detected a single and highly significant locus on Chr 10 (Figure 1) with peak linkage that extends from about 5.7 to 8.9 Mb in males and from 8.9 to 9.6 Mb in females, and maximal –logP score of 10 and higher for males and 9.6 and higher for females (Figure 2) using quantile normalized locomotor data. This strong locus was reported by Philip et al. (2010). But we also now detect a strong second locus for locomotor activation on Chr 16 between 25 and 30 Mb, that only emerges 90 minutes after injection, and that reaches its temporal peak more than two hours after injection (Figure 3). Linkage is significant in both sexes but is several orders of magnitude stronger in females than males—linkages of 10.6 and 4.28, respectively.

**Figure 1.**
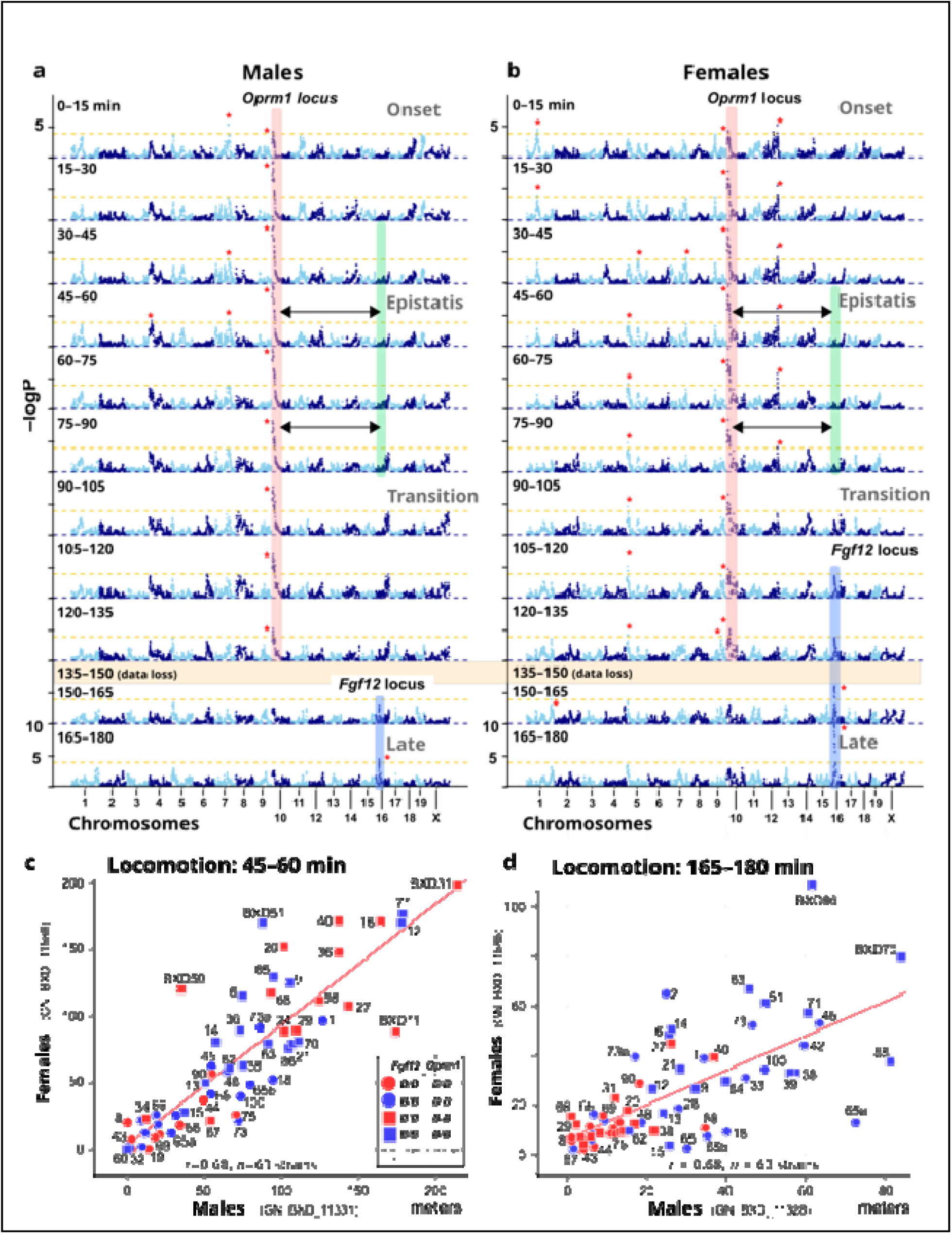
Time series of QTLs for morphine-induced locomotor response

**Figure 2.**
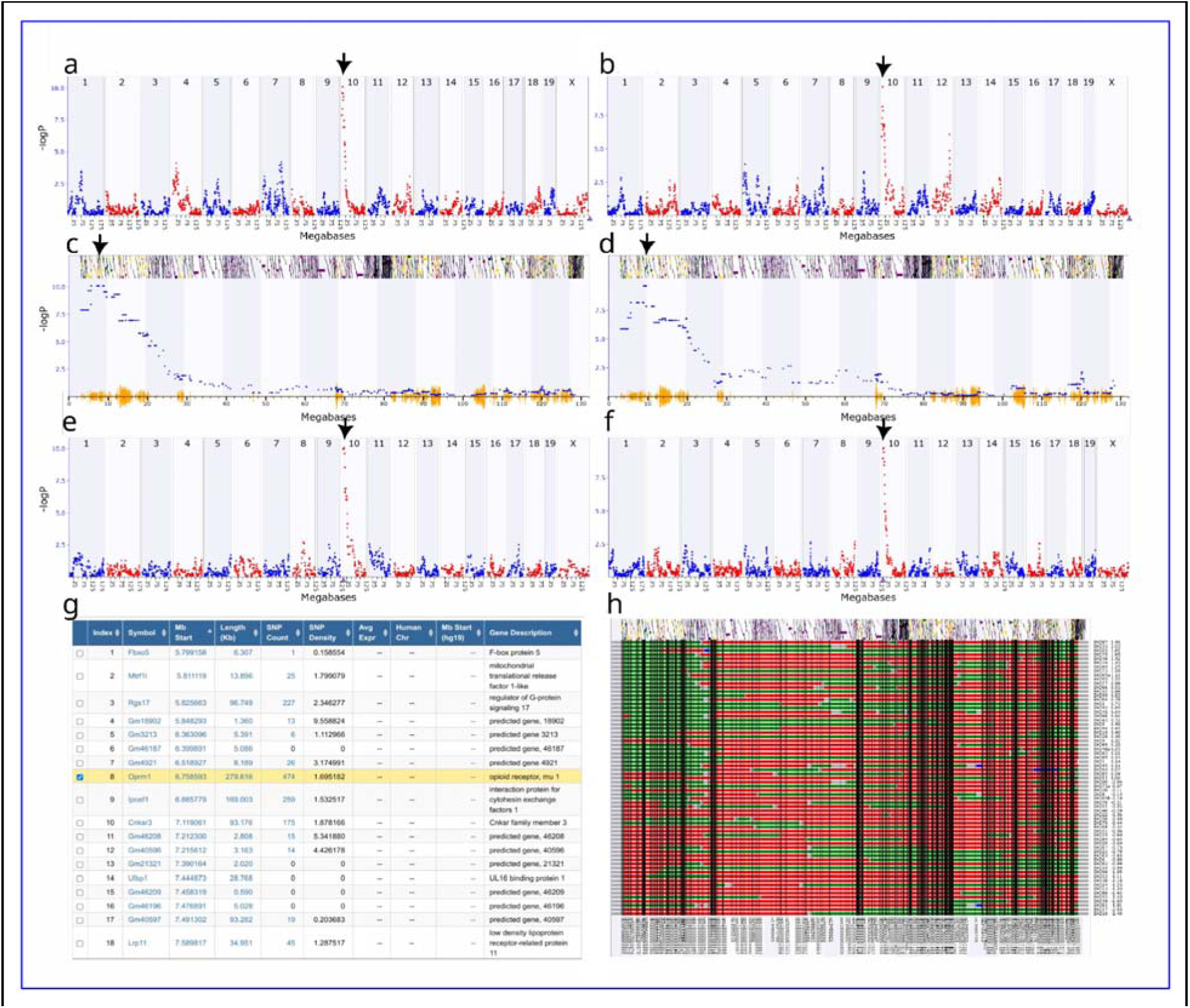
QTLs of morphine-induced locomotion between 45–60 min on Chr 10. (a) QTL for males with a peak –logP of 10.06 (n = 63 strains). (b) QTL for females with a peak –logP of 9.60 (n = 64 strains). (c) Zoomed-in view of QTL in males. (d) Zoomed-in view of QTL in females. (e) Cis-eQTL for Oprm1 in the NAc of the BXDs, with a peak –logP of 10.01 (n = 34 strains). (f) Cis-eQTL for Oprm1 in the hippocampus of the BXDs, with a peak –logP 9.7 (n = 67 strains) on Chr 10 at 5.6Mb. (g) Oprm1 neighborhood in BXD family with SNP densities. (h) Haplotype map of the eQTL region. The “B” of BXD is the mother, and “D” is the father. GEMMA with LOCO was used for all association mapping.

**Figure 3:**
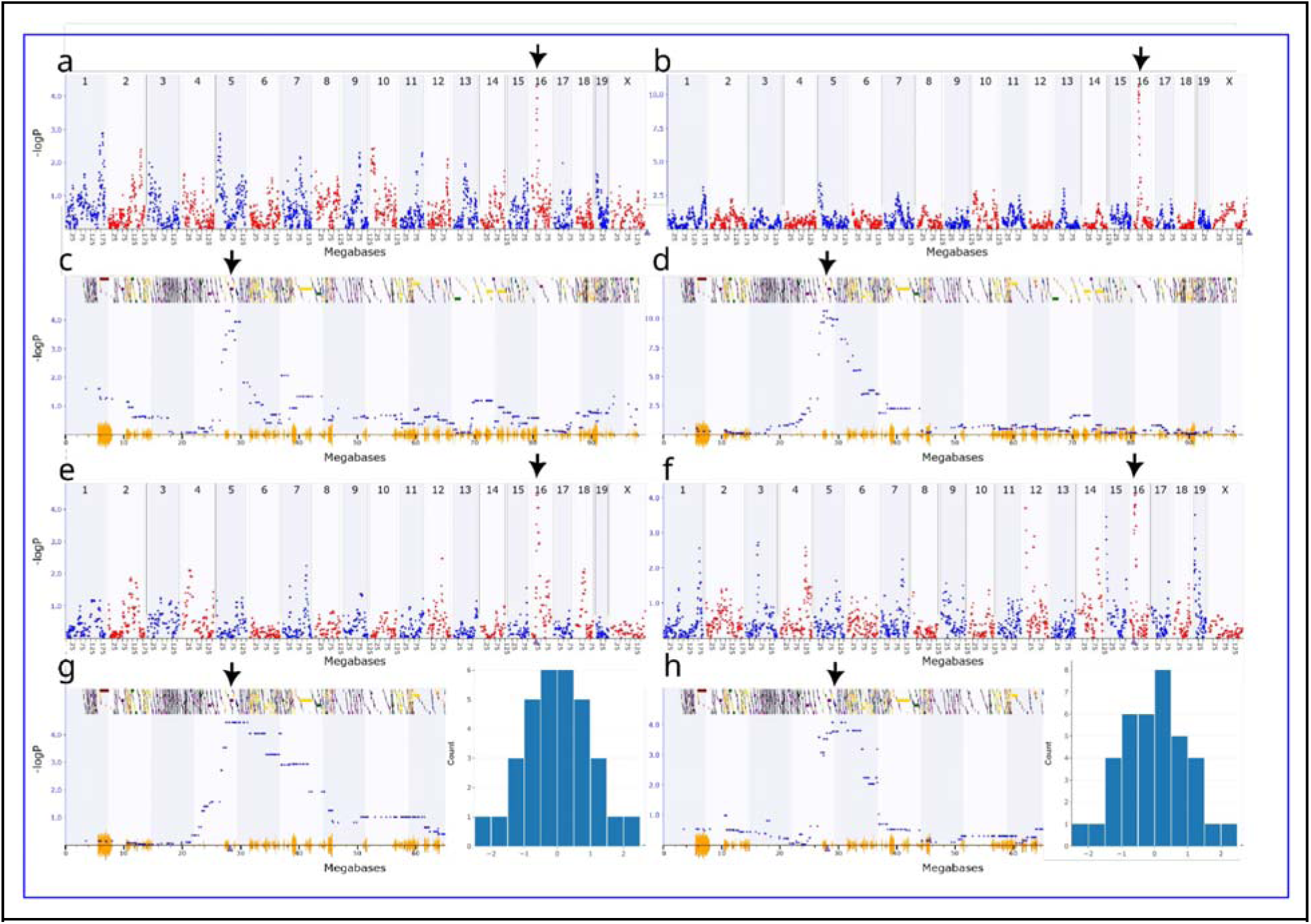
QTLs of morphine-induced locomotion between 165–180 min on Chr 16. (a) QTL in males with a peak –logP of 4.28 (n = 63 strains). (b) QTL for females with a peak –logP of 10.56 (n = 64 strains). (c) Zoomed-in view of the QTL in males. (d) Zoomed-in view of the QTL in females. (e) Cis-eQTL in the striatum of the BXDs, with a peak –logP of 4.43. (f) Cis-eQTL in the VTA of the BXDs, with a peak –logP of 4.06. (g) Histogram of the normalized expression of Fgf12 in striatum and zoomed in view of the QTL region. (h) Histogram of the normalized expression of Fgf12 in VTA and zoomed in view of the QTL region. GEMMA with LOCO was used for all association mapping.

Males and females display roughly similar patterns of linkage on both Chr 10 and Chr 16.

This is consistent with the generally strong positive sex correlations in morphine-induced locomotion responses across all 15-minute intervals. Correlation coefficients peak at about 0.84 to 0.88 from 15 to 75 minutes after injection—the phase during which morphine induces the steepest increase in locomotion—from about 29 to 71 m per 15 min bin (Figure 1c-d). This is also the interval that corresponds to the peak linkage in both sexes near *Oprm1*. Thereafter, correlations drop but are still generally above 0.70 through to 120 min. During the last hour correlations are lower—ranging from 0.57 to 0.68. This drop in correlation corresponds to the period during which effects of the *Fgf12* locus are strongest, and conversely, effects of the *Oprm1* locus are weakest. The distance traveled in the final interval (165 to 180 min) is down to a level slightly lower than that in the first interval—about 25 meters—and the final correlation between sexes is 0.68. This is the interval during which the *Fgf12* locus has a remarkably high linkage of 9.7 and 5.3 in females and males, respectively. In contrast, at this late stage *Oprm1* has linkage below 2.5 in both sexes.

### Candidate gene identification for morphine-induced locomotor response

The mu 1 opioid receptor protein (MOR or OPRM1) is expressed in multiple brain regions and is involved in opioid-induced reward and locomotor response (Contet et al., 2004). In mice the very large *Oprm1* gene is on Chr 10 between 6.76 and 7.04 Mb, just proximal to the linkage peak (Figure 2c-d, GRCm38 assembly). In the BXD family there are over 40 known SNPs, indels, and larger variants segregating in this gene locus, but to the best of our knowledge none alter protein sequence. Given its location just proximal to the QTL peak, its large size, and the high level of genetic variation, *Oprm1* is an uncontroversial candidate for differences in acute morphine locomotor activation (Table 2). But there is some additional support—expression of *Oprm1* mRNA has been quantified in many brain regions across the BXD family, and variants in and around this gene clearly modulate its expression most strongly in nucleus accumbens (NAc) and hippocampus (cis-eQTLs with –logP values >7.0, see examples in Figure 2e,f).

**Table 2.**
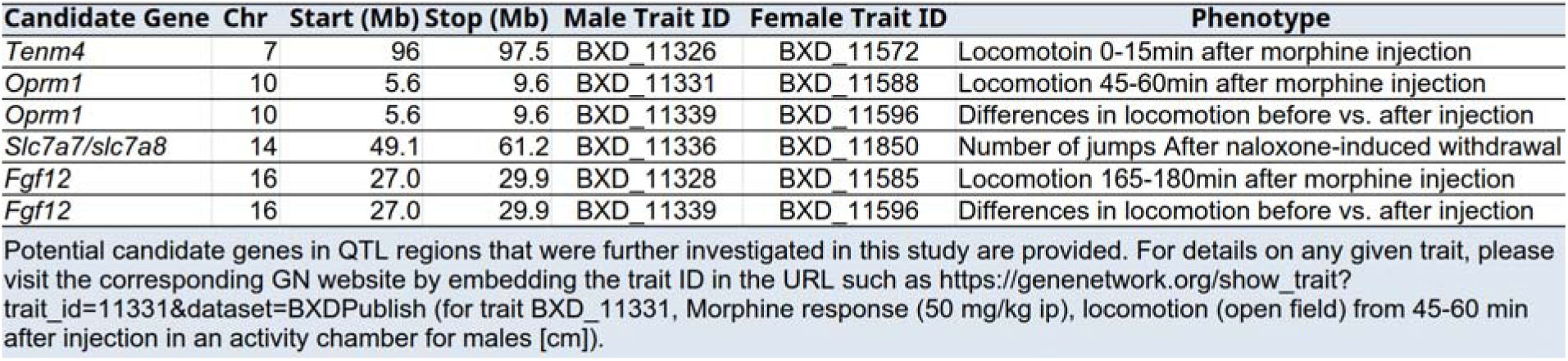
List of Candidate Genes.

Variation in the expression levels of *Oprm1* from the *B* and *D* haplotypes could contribute causally to the Chr 10 locus. Of functional importance, the *D* allele has roughly 50% higher expression in both NAc (Figure 2e) and hippocampus (Figure 2f), an interesting observation given that BXD strains that inherit the *B* allele tend to have a much stronger initial locomotor response.

We restricted our analysis of candidate genes for the novel Chr 16 locus to a 1.5 –logP confidence interval (corresponding to a 95% confidence interval) extending from 27 to 29 Mb (Figure 1). This small region contains only five protein-coding genes and one gene model (*Gm10823*) in the following proximal-distal order: *Ostn, Uts2d*, *Ccdc50*, *Gm10823*, *Fgf12*, and *Mb21d2* (Figure 4). *Fgf12*, previously known as *Fhf1*, is the largest protein-coding gene in this interval (∼600 Kb), with a promoter located close to *Mb21d2* (Figures 3c–d, Figure 4). This is also the strongest biological candidate gene (Table 2), and is already known to interact with the C-terminal region of three sodium channel proteins— SCN9A (Wildburger et al., 2015), SCN8A (Liu et al., 2003), and SCN2A (Wildburger et al., 2015). *Fgf12* encodes a protein that is a member of the fibroblast growth factor (FGF) family. FGFs are involved in response to alcohol (Even-Chen et al., 2017) and morphine (Flores et al., 2010) in key brain reward regions.

**Figure 4:**
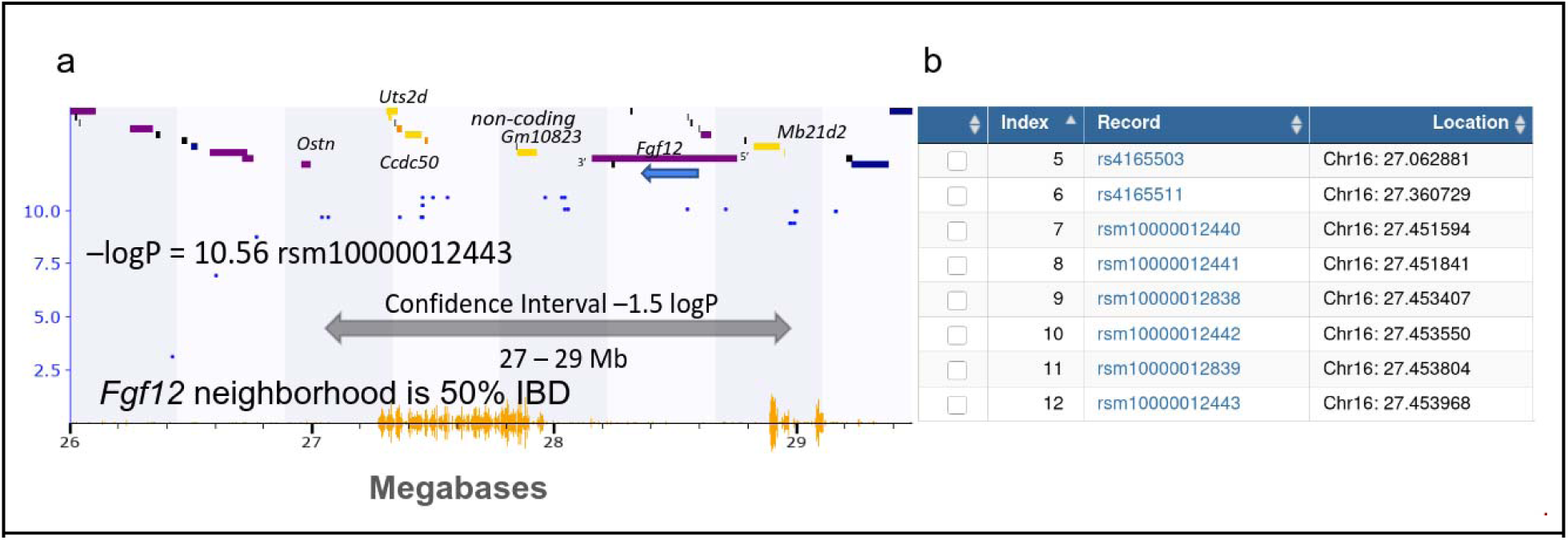
Chr 16 locus at 165-180 min after morphine injection. (a) Zoomed in view of the Chr 16 locus, from 26-29 Mb, showing individual SNPs (blue dots) and genes (purple horizontal lines) in the region. (b) A screenshot of GeneNetwork showing SNPs that inhabit this region. Almost all B vs D SNPs (orange hash along x-axis) are restricted to two regions.

The majority of the *Fgf12* gene is situated in a region that is almost identical-by-descent between the *B* and *D* parental haplotypes. This is highlighted in Figure 4 as the very low SNP density. There are 12 known non-coding variants between *B* and *D* haplotypes, and no known coding variants, however, there are very significant cis-acting expression QTLs associated with *Fgf12* expression differences in the striatum (Figure 3e,g, –logP of 4.4) and the ventral tegmental area (Figure 3f, h –logP of 4.1). The *B* allele is associated with expression that is as much as 60% higher than that of the *D* allele. We identified no coding variants for the other genes in this interval, and *Fgf12* showed the most robust cis-eQTL evidence.

### An epistatic interaction between *Oprm1* and *Fgf12* loci modulates locomotor response to morphine

We detect large additive effects for the locomotor responses to morphine near *Oprm1* at early time points (0–90 min) and near to *Fgf12* at late time points (+90 min). We tested for a possible epistatic interaction between these distinct loci and discovered a strong but transient interaction in both sexes (Figure 5). The epistasis is strongest in the 45–60 min period—precisely at the midpoint between the early and additive *Oprm1* effect and the late and additive *Fgf12* effect (Fig 8c-g, e.g., male trait BXD_11331). Those BXDs that inherit the *B* haplotype of *Oprm1* (defined as rs29339674, 6.7 Mb) as well as the *D* haplotype of the *Fgf12* locus (defined as rs4169220 at 31.6 Mb) have unexpectedly high levels of locomotion compared to all of the other two-locus combinations (Figure 5, Supplementary Figure 3). The –logP value of the epistatic component is in the range of 2.5-4.8 (MF = trait BXD_11845, –logP = 4.2; F trait = BXD_11588, –logP = 4.8; M trait = BXD_11331, –logP = 2.5) and is significant at p < 0.05. The full model that includes this interaction term, as well as both additive effects at *Oprm1* and at *Fgf12* has a remarkably high –logP value of 11.9-13.2 (MF = 13.2, F = 11.9, M = 12.0). This epistatic interaction is preserved as the time window moves forward. For example, maps of the 60–75 min interval (trait BXD_18846) has an epistatic effect with a –logP of 3.0; an additive effect near *Oprm1* with a –logP of 8.2; and an additive effect at *Fgf12* of merely 0.14. The full model in this interval still has a very high cumulative –logP of 11.4. By 75–90 minutes (trait BXD_18847) the interaction effect has dropped to a –logP of 2.2 and the model has a cumulative –logP of 9.40. In marked contrast, over the two intervals from 90 *to* 120 minutes we do not detect significant epistasis. At best, the interaction –logP is only in the range of 1.5–2.0. In the 120–135 and 150–165 minute intervals the originally powerful *Oprm1* additive effect has faded to 2.74 and 1.15, respectively. There is no detectable interaction with the *Fgf12* locus. However, the purely additive effect at *Fgf12* is now much stronger and is detectable for the first time as an independent additive effect with –logP values of 2.1 and 4.2, respectively. Finally, in the last interval—165–180 minutes (trait BXD_11585)—we detect no linkage to the *Oprm1* locus, or any epistasis, but we again pick up a highly significant additive effect at the *Fgf12* locus with a peak –logP that is 4.9.

**Figure 5:**
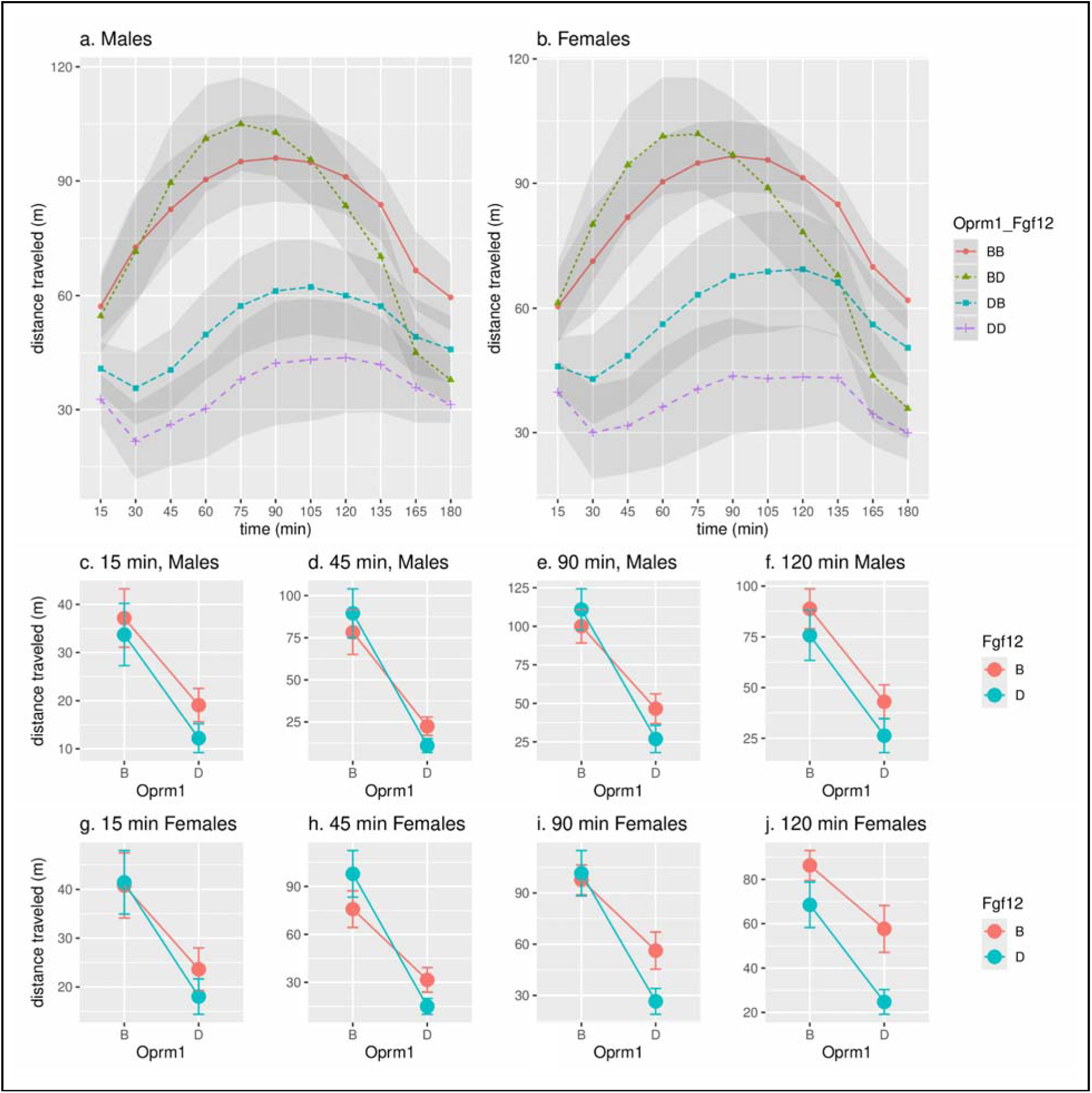
Epistatic Interaction Among Genotype Combinations. (a) Males and (b) females with different combinations of the “B” (i.e., B/B) or “D” (i.e. D/D) genotypes yield different distances traveled after morphine injection over 120 mins. Distance traveled (m) between the two genotypes for each loci is shown for each timepoint in (c-f) males and (g-j) females.

**Figure 6.**
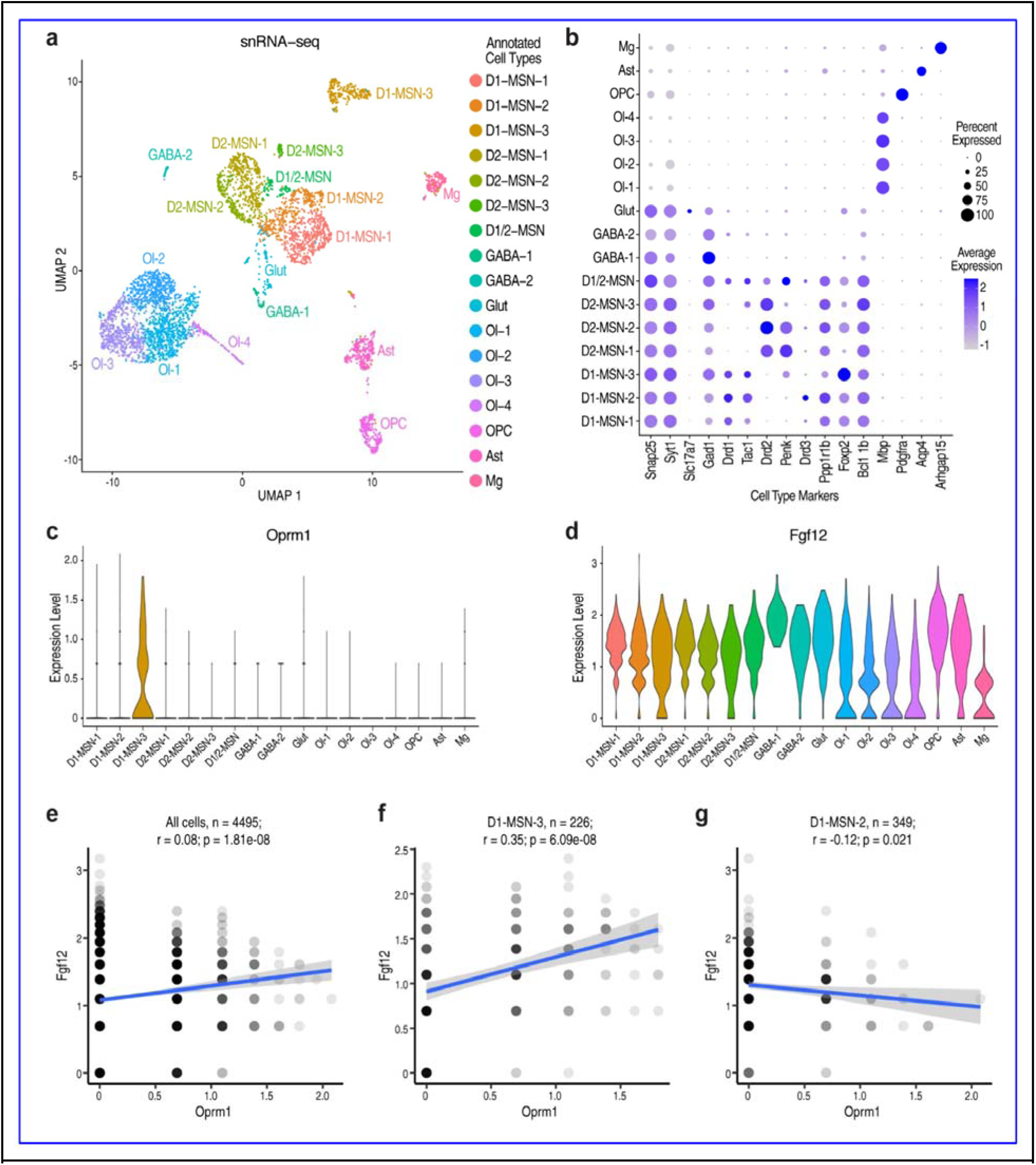
Oprm1 and Fgf12 are positively correlated in rat NAc suggested by snRNA-seq. (a) UMAP visualization of cell clusters from 4495 nuclei from rat nucleus accumbens core. (b) Dot plot showing the expression level of cell type marker genes in cell clusters. The shade of dots denotes normalized and scaled average expression and the size of dots denotes the percentage of cells expressing the gene in each cell cluster. (c-d) Violin plots indicating the normalized and scaled expression level of Oprm1 (c) and Fgf12 (d) across cell clusters. (e-g) Scatter plots showing the correlation relationship between Oprm1

### Cell-type-specific co-expression of *Oprm1* and *Fgf12* in the rat nucleus accumbens core

To further examine whether *Oprm1* and *Fgf12* were co-expressed in the same cells of the NAc, we performed snRNA-seq using a droplet-based approach (10X Genomics). Nuclei were isolated from microdissected NAc cores obtained from two female heterogeneous stock (HS) rats (Carrette et al., 2021). The NAc was chosen due to its central role in opioid reward and the observed strain differences in morphine-induced locomotion (Velásquez et al., 2019). After filtering out low-quality nuclei and potential doublets, a total of 4,495 high-quality nuclei transcriptomes remained with a median number of 3,363 transcripts (unique molecular identifiers) and 1,861 genes detected per nucleus. We then performed SCT transformation (Hafemeister and Satija, 2019), dimensional reduction, and clustering using Seurat (Stuart et al., 2019) and identified 17 cell-type clusters (Figure 6a).

We annotated the cell clusters based on the expression level of established marker genes of the major cell types: neurons (*Snap25, Syt1*), excitatory neurons (*Slc17a7*), inhibitory neurons (*Gad1*), dopaminergic neurons (*Ppp1r1b, Foxp2, Blc11b*), oligodendrocytes (*Mbp*), oligodendrocyte precursor cells (*Pdgfra*), astrocytes (*Aqp4*), and microglia (*Arhgap15*). In addition, we identified 7 subtypes of dopaminergic-receptor neurons. We used expression of *Drd1* and *Tac1* to define D1-type medium spiny neurons (D1-MSNs). The expression of *Drd2* and *Penk* was used to define D2-type medium spiny neurons (D2-MSNs) (Figure 6b, n = 1,768 MSNs total, see Supplementary Figure 4f). MSNs represented the majority (91.8%) of neuronal cells, as expected by previous knowledge of the NAc cellular composition.

*Oprm1* was uniquely expressed in the D1-MSN-3 subtype, while *Fgf12* was expressed across all cell populations (Figure 6c,d). We then performed Pearson’s correlation analysis to test the idea that the epistatic interaction between *Oprm1* and *Fgf12* is mediated by a specific cell type. While a weak but significant positive correlation (*r* = 0.08, *p* = 1.8e-8) between the expression of *Oprm1* and *Fgf12* (Figure 6e) were detected when all cell were included, a much stronger correlation (*r* = 0.35, *p* = 6.1e-8, Figure 6f) was found in D1-MSN-3 cells. In contrast, D1-MSN-2 had a significant weak negative correlation (*r* = –0.12, *p* = 0.02, Figure 6g). In conclusion, D1-MSN-3 is the only cell cluster expressing a high level of *Oprm1* in rat NAc, in which the expression levels of *Oprm1* and *Fgf12* are significantly and positively correlated.

### Constructing and testing Bayesian networks

We hypothesize that genetic variants in the Chr 10 and Chr 16 QTL regions are driving differential expression of *Oprm1* and *Fgf12* in the BXD family and thereby modulating morphine-induced locomotor responses. We used the literature to guide our selection of mediators acting between *Oprm1* and *Fgf12* (Bhushan et al., 2017; Buchsbaum et al., 2002). For example, *FGF12* (Sochacka et al., 2020) and *MAPK8IP2* are linked to a few sodium channel components—SCN2A (Nav1.2) and SCN8A (Nav1.6)(Schoorlemmer and Goldfarb, 2001; Seiffert et al., 2022)—as part of a membrane-associated scaffold concentrated at the axon hillock. The *MAPK8IP2* scaffold protein is thought to modulate JUN amino-terminal kinase signaling, and also interacts with and controls the activity of MAPK8/JNK1 and MAP2K7/MKK7 (O’Leary et al., 2016).

We tested *Scn2a, Scn8a, Map3k11*, *Map3k12*, and *Mapk8ip2* as candidate mediators of the *Oprm1* and *Fgf12* loci and variation in locomotor activity. Expression levels of these transcripts in striatum, VTA, and NAc of BXD strains were passed from GeneNetwork into the Bayesian Network (BN) Webserver (Ziebarth and Cui, 2017) to define the most probable network structure. We constrained the nodes into four tiers: Tier 1 contains the two loci as instrumental variables; tier 2 contains expression estimates of the two prime candidate genes, *Oprm1* and *Fgf12*; tier 3 contains the sodium channels and MAP kinases that potentially mediate both additive and epistatic effects on locomotion; and tier 4 contains the locomotor outcomes expressed in four time-series bins.

The most strongly supported model (Figure 7) recapitulated the cis-eQTL results by indicating that the Chr 10 and Chr 16 loci control the expression of *Oprm1* and *Fgf12* mRNA.

**Figure 7:**
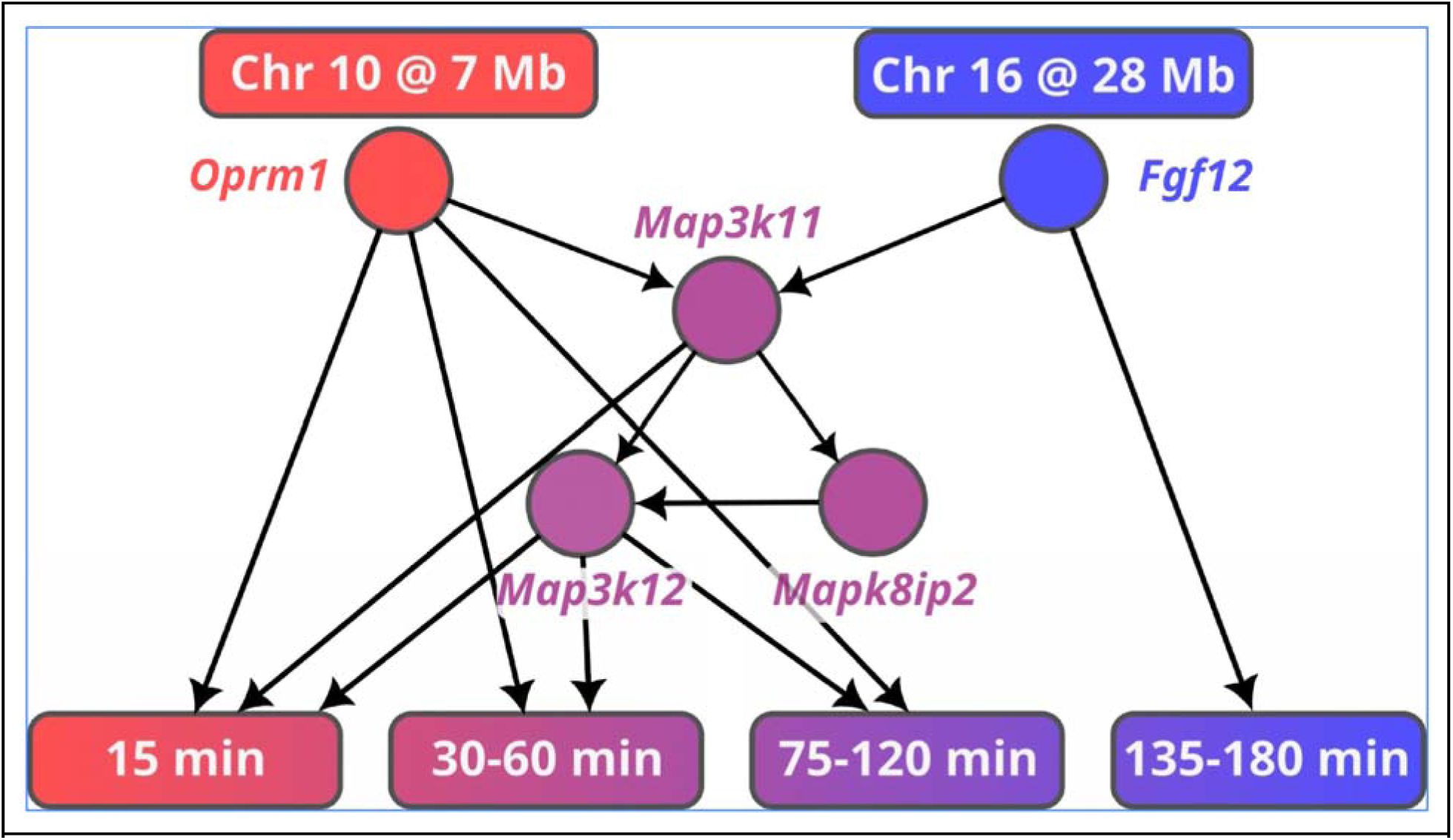
Modeling the mechanistic interactions between genetic variants and phenotypes using Bayesian network. This network illustrates the relationship between genetic variants in the Chr 10 and Chr 16 QTL regions and the differential expression of candidate genes Oprm1 and Fgf12 in the BXD family. Additional gene expression data of MAP kinases were retrieved from Genenetwork.org. Morphine induced locomotion responses at different time bins are also included in the network. This causal hypothesis was developed using the Bayesian network framework available in Genenetwork, where the arrow is indicative of the direction of the causal relationship. Additional genes included in the input of the network construction but had no connection to the final network were excluded in the illustration.

**Figure 8:**
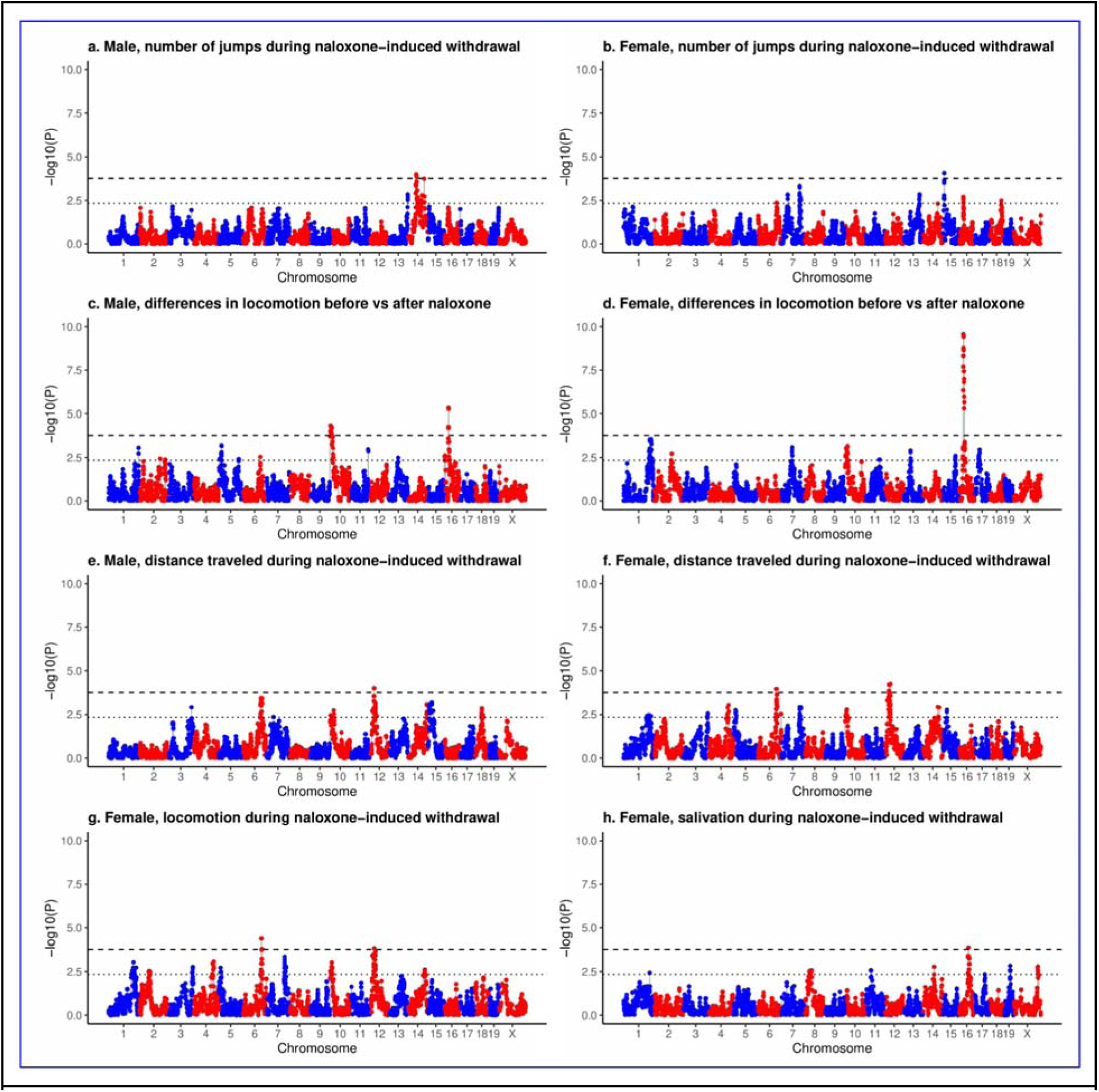
QTLs for naloxone-induced morphine withdrawal responses in males and female BXD mice. Behavior data were quantile normalized and mapped against WGS-based genotypes using GEMMA in Genenetwork.org. (a) Number of jumps 15 minutes after naloxone injection in males. (b) Number of jumps 15 minutes after naloxone injection in females. (c) Change in locomotion measured by the last 15 min of morphine locomotion response minus the first 15 min after naloxone injection in males, with a peak on Chr 10 and a peak on Chr 16 (d) Change in locomotion measured by the last 15 min of morphine locomotion response minus the first 15 min after naloxone injection in females, with a peak on Chr 16. (e) Horizontal activity (distance traveled) 0–15 min after naloxone injection for males. (f) Horizontal activity (distance traveled) 0–15 min after naloxone injection for females, with a peak on Chr 6, and a peak on Chr 12. (g) Number of beam breaks in an open field 0–15 min after naloxone injection in females, with a peak on Chr 6 and a peak on Chr 12. (h) Salivation level in females. Dashed lines: genome wide significance threshold. Dotted lines: threshold for suggestive significance.

This model also supports the causal role of *Oprm1* and its downstream MAP kinases on locomotor response during the first 120 minutes after injection. In contrast, the last 45 minutes are dominated by a direct *Fgf12* effect. Further, *Oprm1* and *Fgf12* directly modulate *Map3k11*, which in turn modulates *Mapk8ip2* and *Map3k12* mRNA expression and regulate locomotion from 0 to 75 min. Further, Mapk8ip2 is embedded in this network with multiple connections.

This Bayesian network aligns well with whole-brain proteomics data (supplementary figure 6). In agreement with the mRNA data, *Oprm1* and *Fgf12* are expressed with opposite polarities, in which the *B* allele is associated with high expression of *Oprm1* and low expression of *Fgf12* and vice versa for the *D* allele. The three MAP kinases are also expressed in higher levels in animals that inherit *B* alleles than *D* alleles.

### QTL mapping of naloxone-induced withdrawal responses

Once the locomotion test was completed (180 min after the morphine injection), naloxone was injected (30 mg/kg in isotonic saline at a volume of 10 ml/kg) to induce a morphine withdrawal response (Philip et al., 2010). Locomotion and other behavioral measurements were taken for both males and females. QTL analysis of these traits were not included in Philip et al. (2010). There are an impressive 16 genome-wide significant QTLs among 5 naloxone-induced withdrawal behavioral responses (Figure 8, Supplementary Table 1).

Figure 8c-d displays the differences in locomotion before and after naloxone injection, with a QTL peak on Chr 10 and 16 that overlap the morphine locomotor QTLs in Figures 2 and 3 and that are likely to correspond to variants in or near to *Oprm1* and *Fgf12*. Our results show that the genotype effect on locomotion was similar before and after naloxone injection (Supplementary Figure 7). We evaluated the biological function of naloxone candidate genes using GeneCup (Gunturkun et al., 2022). Candidates for the locus on Chr 14 for naloxone-induced jumps include *Slc7a7* and *Slc7a8* (Table 2). Both are transmembrane amino acid transporter proteins (Hu et al., 2020) that dimerize with *SLC3A2(Hurkmans et al., 2022; Torrents et al., 1999, 1998)*. Of relevance, *SLC7A8* (LAT2) is an L-DOPA transporter(Crocco et al., 2024).

### Integrating murine data with human GWAS results

There is strong support for *OPRM1* in human OUD GWAS. *OPRM1* variant rs1799971 has been associated with OUD in several studies [e.g., p = 8 x 10^-10^ in Zhou et. al. (2020), p = 2 x 10^-8^ in Hatoum et. al. (2022a), and p = 4.92 × 10^-9^, OR = 1.046, in Deek et. al. (2022)]. A human GWAS of opioid cessation reported linkage to the FGF signaling pathway in which the effect of *FGF12* was nominally significant (*p* = 0.0015, odd ratio = 1.24) (Cox et al., 2020). In gene-based analyses of OUD by Deak et al. (2022), there was a nominally significant signal for *FGF12* (*p* = 0.006), with the top FGF12 SNP rs1553460 also reached nominal significance (OR = 1.015, p = 0.021).

Considering the epistatic interaction between *OPRM1* and *FGF12* and the complexity of human genetic signals (many genes of small effects) it is possible that these two genes function in a network that contains many other genes. We thus developed a list of 500 genes that overlap with *FGF12* and *OPRM1* in the GTEx sample (GTEx Consortium, 2013) (GTEx v8). We then examined whether there is enrichment of members of this network using MAGMA (de Leeuw et al., 2015) in the GWAS by Zhou et al.(2020). Our initial analysis was performed for NAc, putamen, caudate, and adipose tissue for the *FGF12* lists (the latter as a negative control).

For OUD, no significant enrichment was observed for the original 500-gene lists, likely due to the lists’ large size and the limited statistical power of the SUD GWAS. In the GWAS conducted by Deak et al. (2022), where *FGF12* was nominally significant, we found nominally significant enrichment in two of the brain regions, putamen and caudate. Given the low power of this dataset and the broad initial gene list, we refined our analysis to focus on genes specifically correlated with *OPRM1* and *FGF12* in relevant tissues (see Methods). No enrichment was observed for the gene network in the OUD GWAS (p = 0.493). However, this refined approach revealed nominal enrichment (p = 0.038) for the addiction risk factor (Hatoum et al., 2022b).

## Discussion

We analyzed time-dependent behavioral responses to morphine and naloxone collected from the BXD family of mice (Philip et al., 2010) using WGS-based genetic markers and linear mixed models. We discovered a novel association on a Chr 16 locus that overlaps *Fgf12* in both sexes, in addition to confirming an association between locomotor response and a region on Chr 10 that overlaps *Oprm1*. Further, these two loci had a significant but transient epistatic interaction between 45–90 min after morphine injection. According to our transcriptomic data from NAc in rats, *Oprm1* and *Fgf12* are colocalized in a specific subtype of D1-MSN and their expression levels are positively correlated. Analysis of both *OPRM1* and *FGF12* in human GWAS data demonstrated an enrichment of signals associated with SUD phenotypes, and a modest corroboration of variants in the *FGF12* locus on Chr 3q28.

### Additive effect of the *Oprm1* locus

The proximal Chr10 locus associated with early phase locomotor response to morphine contains the *Oprm1* gene. This locus has been associated with the antinociceptive effect of morphine (Bergeson et al., 2001). *Oprm1* encodes the primary receptor responsible for morphine-induced locomotor response (Severino et al., 2020; Smith et al., 2009), and is also responsible for its addiction liability (Ballester et al., 2022; Zhang et al., 2020). Notably, the *Oprm1* mRNA cis-eQTL in the NAc and hippocampus precisely overlaps with the *Mor10a* QTL interval (Figure 2e, 2f), providing strong secondary evidence for *Oprm1* as the causal gene.

Morphine-induced locomotion was affected by *Oprm1* variants (Popova et al., 2019). For example, in the *Oprm1* A112G knock-in mouse, morphine-induced hyperactivity was blunted (Mague et al., 2009). Further, the effect of MOR on locomotion depends on the neuronal cell types involved. For example, when MOR was deleted from D1 neurons, mice had hypolocomotion in response to opioids. By contrast, when MOR was deleted from A2a (adenosine A2a receptor-expressing) neurons, mice displayed increased movement in response to opioids (Severino et al., 2020). These data indicate that *Oprm1* is a strong candidate gene for the Chr 10 locus associated with morphine-induced locomotion response.

BXDs with the *D* allele at the *Oprm1* locus were less sensitive to a 50 mg/kg dose of morphine (Figure 6). The locomotion data demonstrate a blunted motor activity response similar to what was seen in low doses of morphine (5 to 10 mg/kg) (Mori et al., 2004), while BXDs with the *B* allele demonstrate higher sensitivity at the same dose; strains with the *B* allele had an initial biphasic increase in motor activity followed by a drop. This suggests that the *B* allele of *Oprm1* is associated with greater sensitivity or locomotion activity. These data match the differences between the parents of the BXD family. The locomotor response to morphine in C57BL/6J is more pronounced and lasts longer compared to that of DBA/2J (Murphy et al., 2001). This dimorphism may be linked to differences in morphine-induced dopamine release in the NAc in C57BL/6J than in DBA/2J mice (Murphy et al., 2001).

### Additive effect of the *Fgf12* locus

The association between the Chr 16 locus (Figure 1 a,b) and morphine-induced locomotion 150–180 minutes after injection is a novel finding, as is the detection of a time-delimited epistatic interaction of the Chr 16 locus with the *Oprm1* locus. The use of a high-density WGS-based marker set and the LMM (GEMMA) allowed us to detect this and other novel loci not previously detected. Of all positional candidate genes in the Chr 16 locus (Figure 4), *Fgf12* was the most biologically relevant and was supported by strong cis-eQTL in striatum and VTA (Figure 3 e,f). Studies have implicated members of the FGF family, such as *Fgf2* (Even-Chen and Barak, 2019a), FGF receptor 1 (Even-Chen and Barak, 2019b), and FGF receptor 2 (Blackwood et al., 2019) in substance use. It has recently been demonstrated that administration of an *Fgf21* analog in non-human primates decreased alcohol consumption by 50% via an amygdala-striatal circuit (Flippo et al., 2022). Unlike these secreted FGFs, *Fgf12* is an intracellular protein that serves as a cofactor for voltage-gated sodium channels and other molecules and is involved in intracellular signaling events (Schoorlemmer and Goldfarb, 2001). While the full spectrum of its function remains unknown, *Fgf12* has demonstrated a role in locomotion. For example, mice with a null mutation of *Fgf12* have ataxia (Goldfarb et al., 2007). Elevated stress responses have been associated with a reduction in *Fgf12* expression in the prefrontal cortex in rats (McCreary et al., 2016). *FGF12* expression is also elevated in the anterior cingulate cortex of patients with major depressive disorder (Evans et al., 2004).

Variants in human *FGF12* have been linked to cognitive decline (Mohammadnejad et al., 2020), major depression (GENDEP Investigators et al., 2013), schizophrenia (Stertz et al., 2021), and sleep quality (Byrne et al., 2013). Recent evidence also supports the druggability of other intracellular FGFs, such as FGF14 (Dvorak et al., 2025; Singh et al., 2025) and FGF13 (Singh et al., 2025), through their interactions with sodium channels. Our findings suggest that FGF12 could represent a novel therapeutic target for OUD by modulating its interaction with sodium channels.

### The epistasis of morphine locomotor activation

We identified a highly significant epistatic interaction between the *Oprm1* and *Fgf12* loci 45–90 min after morphine injection (Figure 6), that erodes in the following 30 mins, providing strong pharmacological constraints on mechanisms. Those strains that inherit both the *B/B* genotype at *Oprm1* and the *D*/D genotype at *Fgf12* have significantly higher activity levels compared to those strains that inherit the other three genotype combinations. Our mapping extends in intervals of 15 minutes over a 3 hour period. We can divide this interval into four phases (Figure 1).The first phase after the morphine injection is influenced only by DNA variants in the *Oprm1* locus. The second phase— from 45 to 90 minutes—is characterized by the strong but transient epistatic interaction of *Oprm1* and *Fgf12* loci. The third phase is a kind of hiatus that extends from 90 to 120 minutes in which we do not detect any epistatic interactions. Only *Oprm1* is active during this phase. In the final phase—from 120 to 180 minutes—the *Oprm1* effect is exhausted and now the *Fgf12* locus acts independently for the first time with a purely additive effect on activity. The hiatus phase may be explained mechanistically by effects of other loci that we are not yet able to detect reliably. Note for example the transient female-specific locus on Chr 5 that fills part of the hiatus. This complex time course of transitioning in genetic control on morphine induced locomotion is likely caused by molecular cascades triggered by the activation of the *OPRM1* receptor and its signalling molecules and downstream molecular network linked to *FGF12*. To some extent we have tried to bridge the gap between the genetics of morphine responses to possible molecular cascades using Bayesian causal modeling as in Figure 8.

Complex traits, such as OUD, are controlled by many genes that interact with each other and their environment (Crist et al., 2019). While improved statistical approaches (D’Silva et al., 2022; Wei et al., 2014) and novel machine learning methods (Chicco and Faultless, 2021) are starting to enable the study of epistasis in human data, most human genetic studies either did not have the power to detect epistasis or ignored its effect (Carlborg and Haley, 2004; Uffelmann et al., 2021). In contrast, many epistatic interactions have been identified using model organisms (Mackay, 2014), mostly due to the ability to obtain data from individuals with controlled genotypes and measuring phenotypes in well controlled environments (Campbell et al., 2018). Identifying gene-gene interactions in model organisms can provide candidates to be evaluated in the human population for its potential in improving the prediction of individual disease risk and the application of personalized therapies.

### Cell-type specific gene expression in NAc

While gene interaction is nearly always measured statistically without regard to biological mechanism, we explored the utility of snRNA-seq in identifying cellular and molecular mechanisms of this interaction by examining gene expression in NAc, a brain region relevant to the biological effect of morphine. We focused on NAc because differences in morphine-induced dopamine release in NAc correspond with the locomotor response in the parental strains of the BXD mice (Murphy et al., 2001). Further, mice lacking *Drd1* are not responsive to acute cocaine-(Karlsson et al., 2008; Xu et al., 1994) or morphine-(Wang et al., 2015) induced locomotor response. While our analysis identified 17 distinctive cell types (Figure 7a), *Oprm1* was detected almost exclusively in the D1-MSN-3 subtype that have high expression of *Foxp2* and low expression of *Ppp1r1b* (i.e. *Darpp-32*; Figure 7b,c). The total number of genes (n_feature RNA) and other quality control (QC) parameters in D1-MSN-3 are comparable with other neuronal cell types. In contrast, *Fgf12* was expressed in most of the neuronal and glial cell types (Figure 7d). The expression level of *Oprm1* and *Fgf12* were positively correlated (r = 0.36, p = 6e-8) in D1-MSN-3 but not in D1-MSN-2 cells, indicating a potential cell-type specific mechanism for the epistatic interaction.

MSNs in the NAc have been the subject of intense research. A recent snRNA-seq study of the NAc in rat brains also detected this specific subtype expressing high levels of *Oprm1*, which is selectively labeled by the expression of *Chst9* (Andraka et al., 2023; Savell et al., 2020). These data suggest that D1-MSN-3 is a highly likely location for the interaction between *Orpm1* and *Fgf12*. Notably, this cell type is conserved also in primate and human NAc snRNA-seq data (Andraka et al., 2023), suggesting that it may play a critical role in opioid responses across species.

To further confirm our results and allow for analysis of correlations, we queried data from the *Ratlas* (https://day-lab.shinyapps.io/ratlas/), which reveals that *Fgf12* is indeed widely expressed in all cell types in other datasets, and that *Oprm1* was not detected in either *Drd1* or *Drd2* MSN neurons. This is most likely due to the lack of detection of the MSN subclass that is equivalent to the D1-MSN-3 subtype that we identified (Supplementary Figure 5a-d). However, cultured cell data do express *Oprm1* in both *Drd1* and *Drd2* cells (Supplementary Figure 5e-f). While out of the scope of this paper, future work will be needed to assess the impact of the *Oprm1-Fgf12* interaction on the activity of the D1-MSN-3 population in mediating opioid responses.

### Bayesian causal network modeling of *OPRM1* and *FGF12*

We included gene expression data of several MAP kinases and sodium channels with known relationships with *Oprm1* and *Fgf12* as the input for our BN. The resulting network (Figure 8) indicated that *Mapk8ip2*, *Mapk3k11* (*Mlk3*), and *Map3k12* (*Dlk*) form a highly interactive network mediating the interaction between *Oprm1* and *Fgf12* in regulating their effects on morphine-induced locomotor response. In addition to validating the results on genetic influence on *Oprm1* and *Fgf12* expression and morphine-induced locomotor response, the network identified *Map3k11* as a potential nexus that links *Oprm1* and *Fgf12. Map3k11* further activates other MAP kinases such as *Mapk8ip2* and *Map3k12* to modulate locomotion before 120 min. Thus, *Map3k11* is a potential mediator of the epistasis between *Oprm1* and *Fgf12*. Our BN did not contain a direct relationship between *Fgf12* and *Mapk8ip2*, the encoded proteins for which directly bind to each other (Bhushan et al., 2017). It is possible that these two proteins do not regulate each other’s mRNA levels, which was used in our BN. While our modeling suggest that the epistatic interaction between *Oprm1* and *Fgf12* is mediated through *Map3k11*, further research, such as those using Mendelian randomization studies and other forms of mediation analysis, could provide further understanding of the interactions between MAP kinases, *FGF12*, and *OPRM1*.

### Genetic modulation of naloxone effects

Naloxone is a potent opioid antagonist that reverses opioid overdoses (Theriot et al., 2022), and is often used in treatment settings or as a primary overdose prevention method.

Pharmacogenomics has primarily focused on individualized treatment for pain, with less priority on OUD. However, understanding the role genetics plays in naloxone’s mechanism could initiate potentially more personalized and effective treatments for OUD. We identified multiple significant associations between behavioral measures of the effect of naloxone on Chrs 3, 5, 7, 10, 11, 12, 13, 14, and 18. These loci are likely to contain novel genes that play a role in OUD. For example, *Slc7a7* and *Slc7a8* are candidate genes for the number of jumps after naloxone injection. Both of these genes are a part of the solute carrier family, which play a large role in the absorption of drugs and xenobiotics (Lin et al., 2015), and are expressed in the brain.

Similar to the morphine locomotion time series phenotypes, there are significant QTL for the differences in locomotion before and after naloxone injection (Figure 8c-d) on Chrs 10 and 16. Mice with the *B* allele at the *Fgf12* locus still had elevated locomotion during 165-180 min compared to those with the *D* allele. Injection of naloxone eliminated the remaining effect of morphine and brought locomotion in both genotypes close to baseline level (Supplementary Figure 7). The difference in locomotion before vs after naloxone injection thus amplified the net effect of morphine at this late phase and further confirmed that this effect was mediated via the MOR. The highly significant association (–logP = 5.34 in males and –logP = 9.57 in females) between the Chr 16 locus and this phenotype provided a strong confirmation for its role in late phase response to morphine. Analysis of these genes in the homologous human regions in larger populations are needed to translationally confirm and extend the validity of these results.

### Integration with human data

While our attempt to integrate murine data with human GWAS results demonstrated that the pipeline works, the lack of enrichment for the network in OUD GWAS could likely be due to the small sample size in the original human data for that specific SUD. This could also likely be due to the lack of data specification, such as the differences between examining physical dependence vs the development of OUD, initial use vs problematic use, and withdrawal vs relapse (Sanchez-Roige et al., 2021).

In summary, our study confirmed that the *Oprm1* locus strongly modulates the early phase locomotor response to morphine and identified a novel association between *Fgf12* locus and the late phase response to morphine. To the best of our knowledge, the epistatic interaction between the *Oprm1* and *Fgf12* loci during the middle phase of locomotor response is the first demonstration of a transient time-dependent epistatic interaction modulating drug response in mammals—a finding with interesting mechanistic implications. Our snRNA-seq analysis suggests that the D1-MSN-3 subpopulation of dopaminergic-receptor neurons might be critical in the epistatic interaction between *Oprm1* and *Fgf12*. Our study further shows that the interaction between the *Fgf12* and *Oprm1* genes is mediated by *Mapk8ip2, Map3k11*, and *Map3k12*, and that the activity of these proteins is modulated by the presence of *Fgf12* variants. Finally, this work demonstrates how high-quality Findabile, Accessible, Interoperabile, and Reusable (FAIR+) phenotypes can be used with updated data sets to yield striking results, and how joint mouse and human neurogenomic and GWAS results can be merged at gene and network levels for bidirectional validation of SUD variants and molecular networks.

## Methods

### Animal and Behavior Data Collection

Morphine-induced locomotor responses and naloxone-induced withdrawal were recorded for 64 BXD RI strains. All animals were young adults (8–9 weeks) reared at the Oak Ridge National Laboratory in 2007–2008. Detailed methods were reported in the original publication by Philip et al (2010). Briefly, the testing protocols included giving each mouse a single i.p. injection of morphine sulfate (50 mg/kg in isotonic saline at a volume of 10 ml/kg), followed by immediately placing the mice into an activity chamber (43.2 cm L × 43.2 cm W × 30.4 cm H, ENV-515, Med Associates, St Albans, VT, USA). Each chamber contained two sets of 16 photocells placed at 2.5 and 5 cm above the chamber floor. Activity was measured as photocell beam breaks and converted into horizontal distance traveled (cm). This behavior and rearing were recorded for 3 h and were exported as the sum of 15 min bins. Then, each mice in these cases received an injection of naloxone (30 mg/kg in isotonic saline at a volume of 10 ml/kg, i.p.), and were immediately returned to the chambers for an additional 15 minutes. Naloxone’s effects on locomotor activity levels, and signs of withdrawal were recorded for 15 minutes, including the number of jumps, fecal boli, urine puddles, wet dog shakes, instances of abdominal contraction, salivation, ptosis and abnormal posture were recorded for 15 minutes). The number of animals used per strain was 7.8 (mean ± SD) for females and 6.6 for males. Testing occurred at about 8 to 9 weeks after birth.

### QTL Mapping in GeneNetwork

We reanalyzed 105 phenotypes from the Philip et al. (2010) study available in GeneNetwork.org, a database containing phenotypes and genotypes, and also serves as an analysis engine for quantitative trait locus (QTL) mapping, genetic correlations, and phenome-wide association studies (Mulligan et al., 2017; Sloan et al., 2016; Watson and Ashbrook, 2020). The trait IDs and their most significant loci are shown in Supplementary Table 2. The data for mapping QTLs consist of polymorphic genetic markers and quantitative trait values for strain means. Statistical heritability was also evaluated to estimate the degree of variation among the morphine and naloxone traits due to genetic variation. The original analysis (Philip et al., 2010) was performed with the expanded BXD strain to evaluate complementary behavioral phenotyping using Haley-Knott regression in GeneNetwork. In our reanalysis, we first quantile normalized the trait data. We then used the newly implemented GEMMA (Zhou and Stephens, 2012) with the LOCO option, which corrects family structure, to perform genetic mapping using approximately 7,000 markers obtained from whole genome sequencing. Candidate genes were selected from the confidence interval of a one-LOD drop-off from the peak statistical significance as determined previously (Philip et al., 2010). The criterion for genome-wide significance was a –logP value of 3.77 for the BXDs. Loci labeled by genetic markers could act independently at all time points or possibly in an epistatic interaction at a few time points.

Therefore, we also examined epistatic interactions between two loci by performing a pair-scan, implemented in GeneNetwork (v1). This scan generated a matrix map output. Functional enrichment of these networks were obtained using WebGestalt (Zhang et al., 2005). The biological relevance of candidate genes in the context of SUD was queried using Genecup (Gunturkun et al., 2022). We conducted most early exploratory analysis using the web version of Genenetwork. For final analysis, we used the API interface of genenetwork (Mulligan et al., 2017) to conduct standardized analysis and generate uniform figures for all phenotypes of interest.

### Brain samples for snRNA-seq

Brain samples from 2 female HS rats were obtained from the oxycodone tissue repository at UCSD (Carrette et al., 2021). These rats were selected based on their low addiction-linked behavior phenotypes from a larger cohort characterized in the oxycodone self-administration paradigm. The brain tissues were collected from rats euthanized after 4 weeks of prolonged abstinence from oxycodone intravenous self-administration (Carrette et al., 2021). Brain tissue was extracted and snap-frozen (at −30°C). Cryosections of ∼500 μm (Bregma 2.28-0.72 mm) were used to dissect the NAc core punches on a −20°C frozen stage. Punches from 3 sections were combined for each rat.

### snRNA-seq library preparations

snRNA-seq library was performed using the Chromium Next GEM Single Cell Multiome Reagent Kit A (catalog number 1000282) following Chromium Next GEM Single Cell Multiome ATAC + Gene Expression Reagent Kits User Guide (10X Genomics). Approximately 10,000 nuclei were loaded per reaction, targeting recovery of 6,000 nuclei after encapsulation. After the transposition reaction, nuclei were encapsulated and barcoded. Next-generation sequencing libraries were constructed following the User Guide. Final library concentration was assessed by Qubit dsDNA HS Assay Kit (Thermo-Fischer Scientific) and post library QC was performed using Tapestation High Sensitivity D1000 (Agilent) to ensure that fragment sizes were distributed as expected. Final libraries were sequenced using the NovaSeq6000 (Illumina).

### Bioinformatic analysis of snRNA-seq data

Raw sequencing data were converted to FASTQ format using bcl2fastq (Illumina). The FASTQ data were firstly aligned to the Rattus Norvegicus mRatBN7.2/rn7 genome and then aggregated into one sample file using CellRanger version 2.0.0 using default parameters. The output files were analyzed using Seurat version 4.1.1 (Stuart et al., 2019). Nuclei with gene numbers between 500 and 6000, RNA counts between 1000 and 16000, the percentage of mitochondrial gene reads lower than 2%, and the percentage of small 40S or large 60S ribosomal (Rps and Rpl) gene reads lower than 1% were considered as high-quality cells and kept for further analyses. To eliminate potential doublets, we used DoubletFinder 2.0.3 and removed 5% nuclei identified as doublets (McGinnis et al., 2019).

High-quality singlet nuclei were then normalized and scaled using SCT transformation (Hafemeister and Satija, 2019) with the percentage of mitochondrial genes, RNA counts, and sample ID as covariates. The dimensional reduction was performed using PCA and the first 30 PCs were used for KNN graph construction and clustering using the Louvain algorithm. Uniform manifold approximation and projection (UMAP) was used for visualization of the clusters.

The dot plots and violin plots were generated using Seurat (Stuart et al., 2019). Pearson correlation was computed in R 4.2.1 using cor.test and p<0.05 was considered significant.

### Human translational data

We reviewed OUD and other relevant published GWAS data in previous literature with variants in *OPRM1* and *FGF12*. We also extended our review to looking at potential roles of *FGF12* in humans specifically. Then, All the files for *FGF12* and *OPRM1* correlations for each individual brain region as well as some controls were downloaded using GTEx v8 (GTEx Consortium et al., 2017) and GeneNetwork and then extracted out the shared genes with a liberal cut-off of 500 genes. We ran a genome-wide enrichment analysis using MAGMA, a powerful tool that identifies genes and gene-sets associated with a specific disease for analysis (de Leeuw et al., 2015). However, upon running our genome-wide enrichment analysis for OUD, we realized our initial search was too broad and much larger than in previous literature (see results section). To reduce the amount of noise we identified genes that overlap between *FGF12* and *OPRM1* in a given tissue, and identified genes correlated specifically to each *OPRM1* and *FGF12* in several tissues. To do this we developed a list of genes (and Ensembl IDs) in each brain region to examine the number of occurrences each gene is seen across this data. We narrowed down our search to the top genes with high counts and compared them to the same number of genes with a count of only one. The count is the number of times the gene is associated with both *FGF12* and *OPRM1* in all the regions/tissue datasets in GeneNetwork. Association is defined as among the top 500 genes ranked by correlation with *FGF12* or *OPRM1*.

### Bayesian network modeling

The BN server located at https://bnw.genenetwork.org/sourcecodes/home.php was used. Raw data files containing genotypes for the *Oprm1* and *Fgf12* loci, morphine-induced locomotion, and expression levels of relevant genes (sodium channels, MAP kinases) were transferred from Genentwork. We separated our nodes into 4 tiers to label chromosome location, candidate gene, signaling molecules, and phenotype (morphine locomotion) each as a separate tier.

These are used to organize the nodes into different layers. Each tier represents a different layer of complexity. We permitted causal connections to flow from the first tier to the second tier, as well as from the second tier to the third tier. We also allowed interaction between nodes located within the second or the third tiers.

## Data availability

snRNA-seq data has been deposited into NCBI GEO (accession number: GSE214388).

## Supplementary Figures and Tables

**Supplementary Figure 1:**
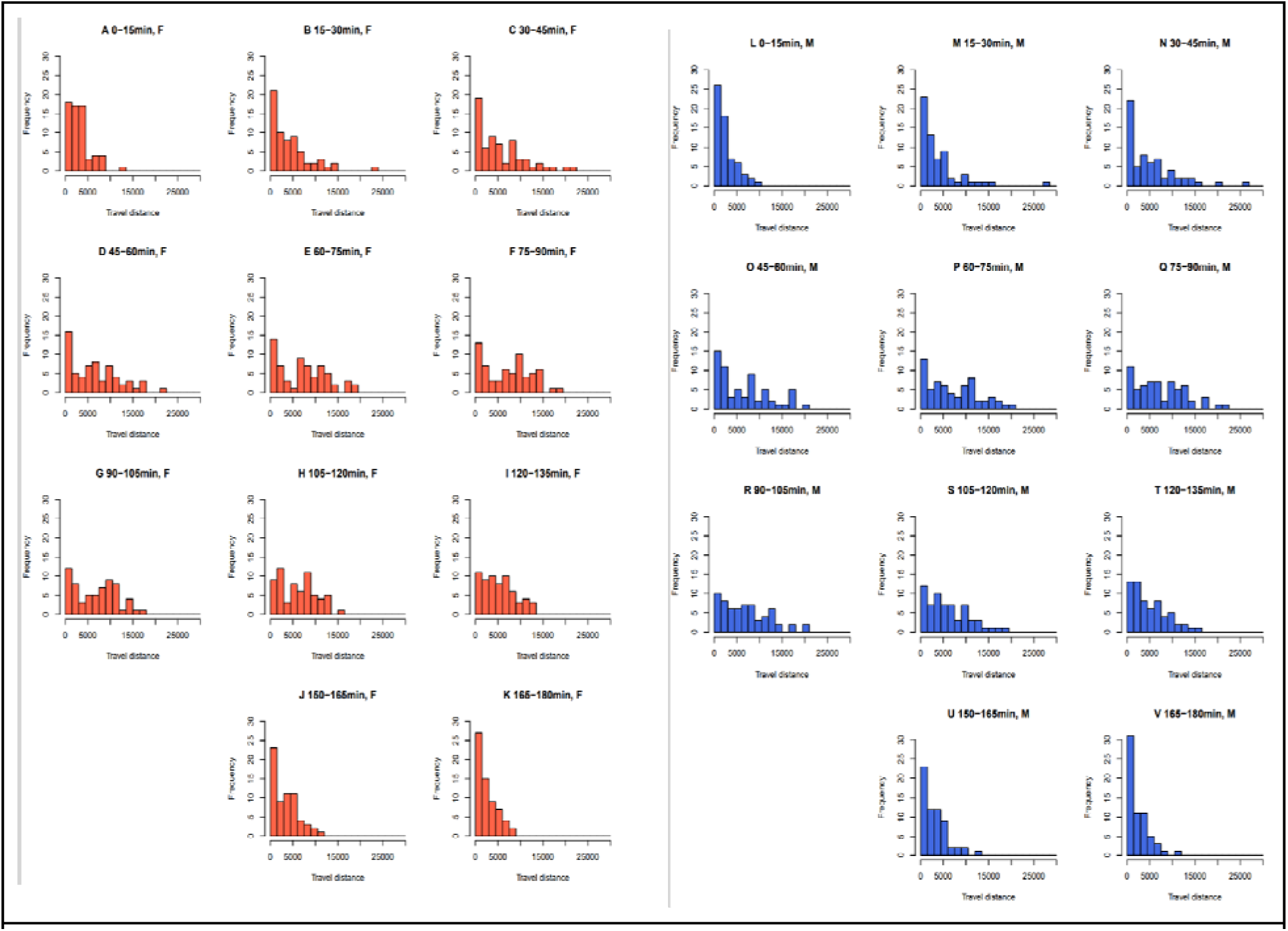
Distribution of raw locomotion data after morphine injection. Distribution of raw data for locomotion after morphine injection at different time intervals in females (orange A-K) and males (blue L-V). Data for 135 to 150 minutes were lost for both sexes.

**Supplementary Figure 2:**
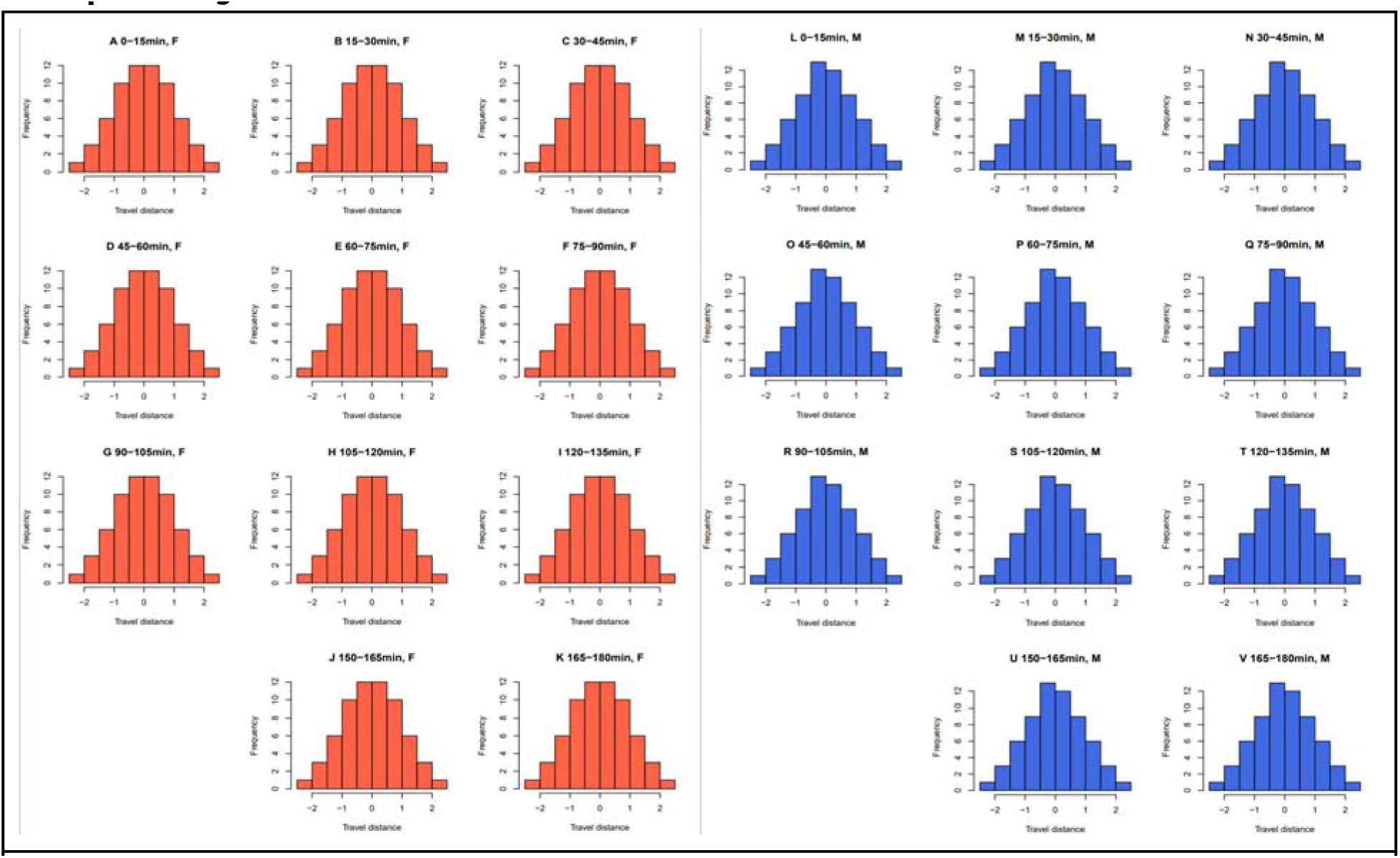
Distribution of quantile-normalized locomotion data after morphine injection. Data are plotted for each time bin for both females (orange A-K) and males (blue L-V). Data for 135 to 150 minutes were lost for both sexes.

**Supplementary Figure 3:**
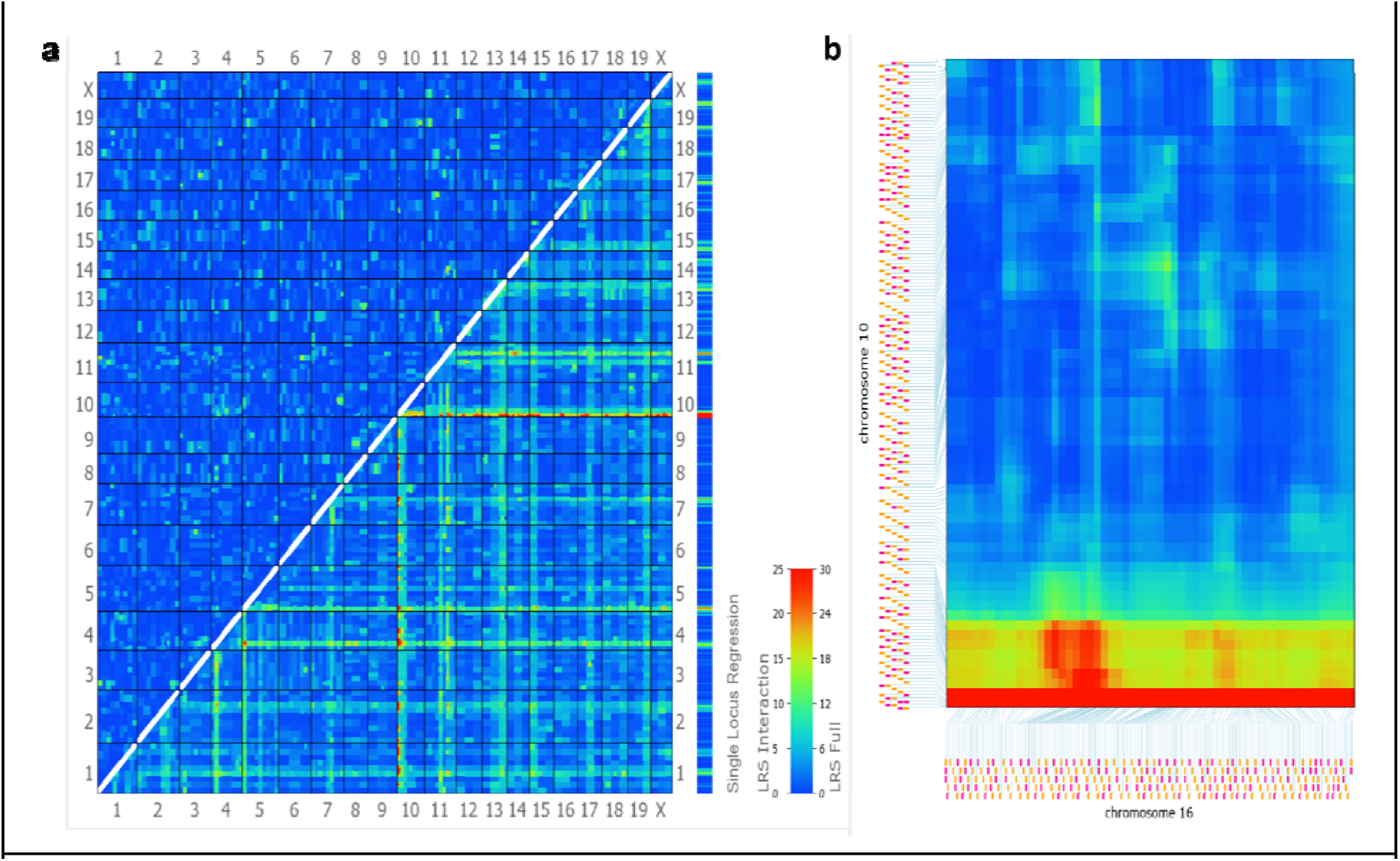
Pairwise linkage statistics across the genome. (a) Genome-wide two-dimensional heat map showing all pairwise marker combinations, illustrating both additive (main) effects and epistatic (interaction) effects. The upper-left triangle displays LOD scores for interaction terms, while the rightmost column shows the main effect (single-locus) genome scan. (b) Zoomed view of linkage statistics between chromosomes 10 and 16. The red band along the bottom represents a strong additive effect on chromosome 16.

**Supplementary Figure 4:**
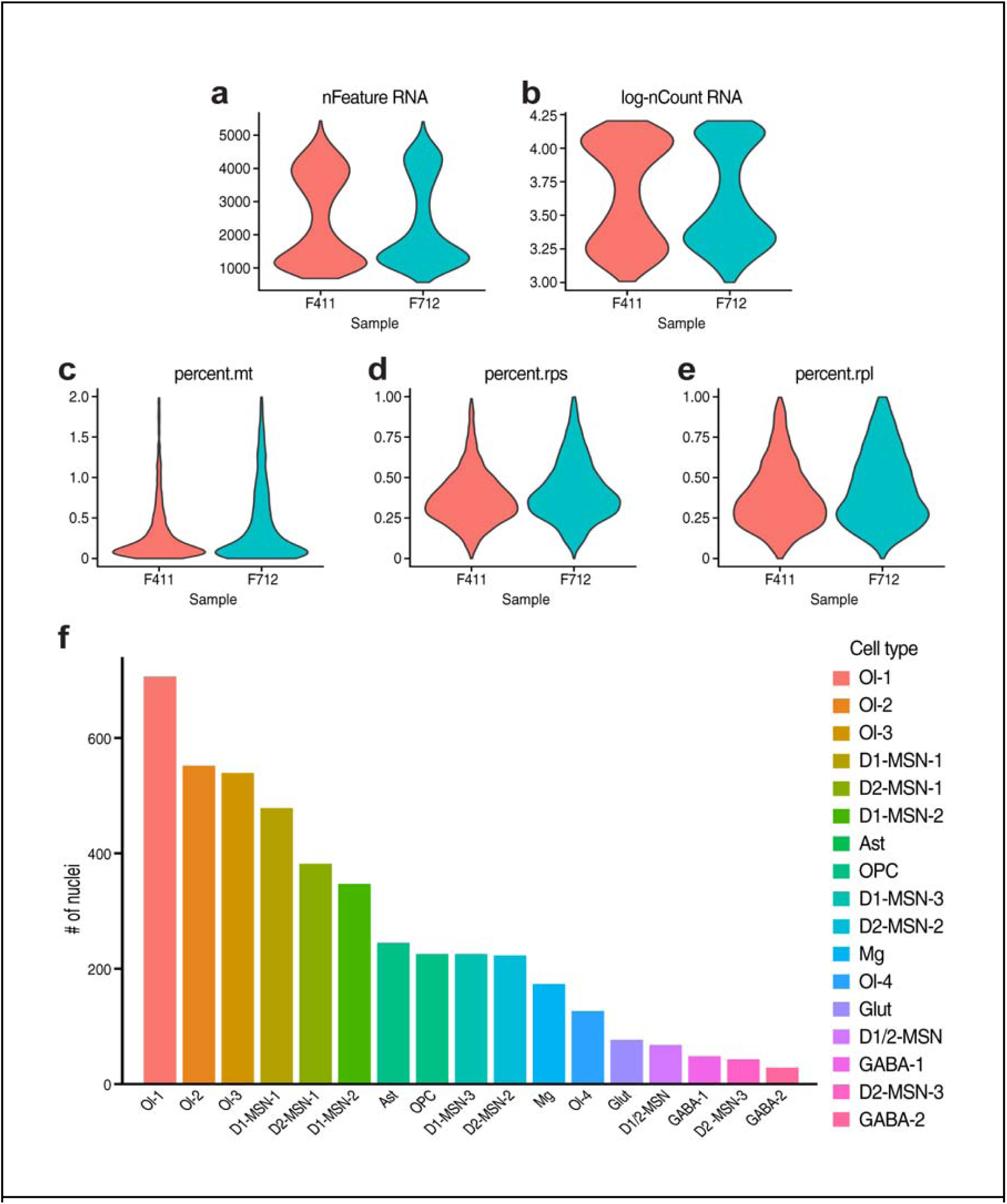
Quality control (QC) of snRNA-seq and number of cells per cluster. (a-e) Violin plots showing QC parameters used for selecting high-quality nuclei in each sample: (a) Unique gene numbers per nuclei (nFeature_RNA). (b) Log-transformed total read numbers per nuclei (log-nCount_RNA). (c) Percentage of mitochondrial gene reads. (d) Percentage of small 40s ribosomal (Rps) gene reads. (e) Percentage of large 60s ribosomal (Rpl) gene reads. (f) Number of nuclei per cell cluster. D1-MSN, Drd1-expressing medium spiny neuron; D2-MSN, Drd2-expressing medium spiny neuron; GABA, GABAergic inhibitory neuron; Glut, glutamatergic excitatory neuron; Ol, oligodendrocyte; OPC, oligodendrocyte precursor cell; Ast, astrocyte, Mg, microglial cells.

**Supplementary Figure 5:**
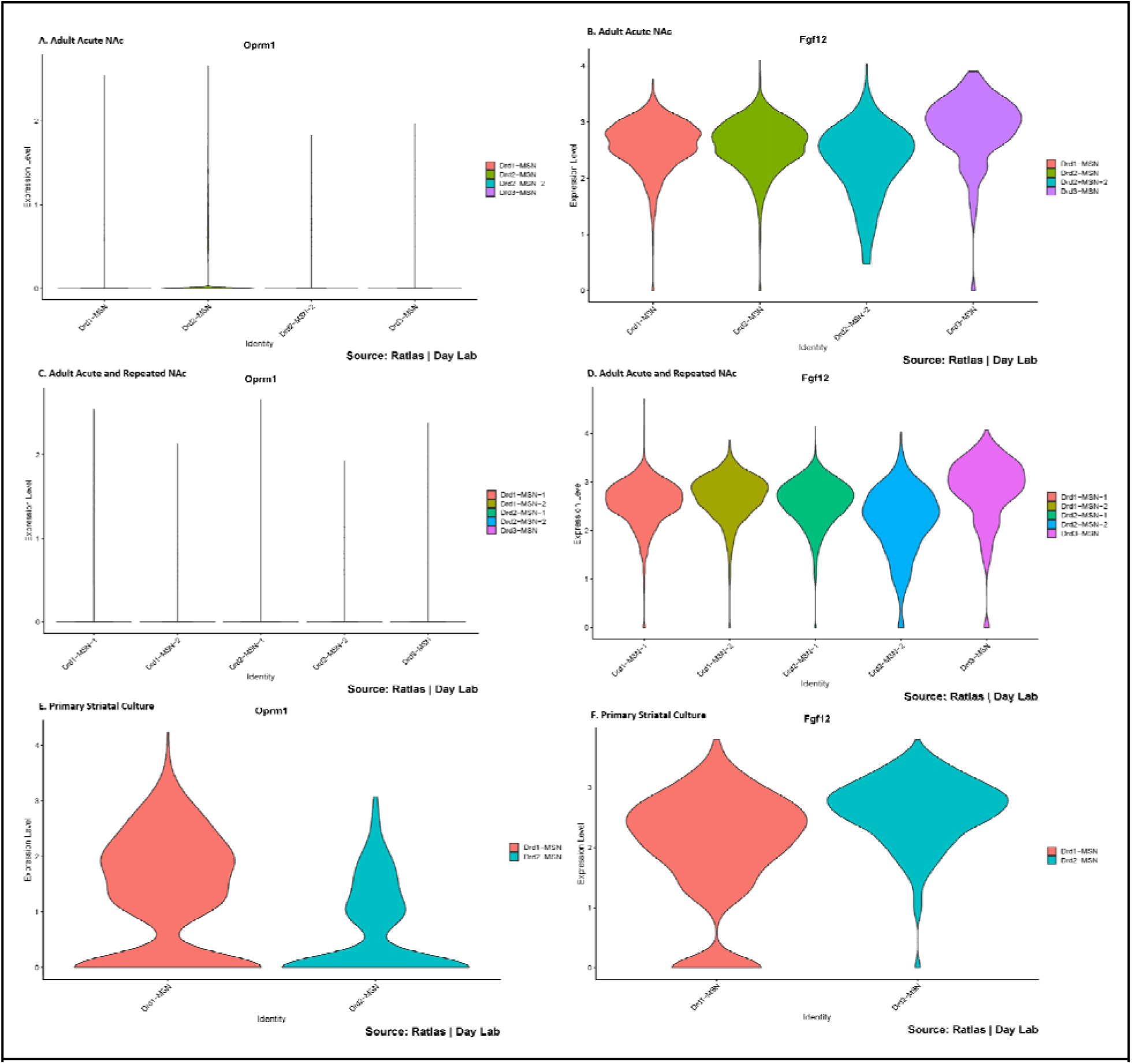
Gene Expression of D1 and D2 Type Medium Spiny Neuron Subtypes. Expression of Oprm1 and Fgf12 in acute and repeated nucleus accumbens and in primary striatal culture from the Ratlas database. (a) Expression of Oprm1 in acute NAc. (b) Expression of Fgf12 in acute NAc. (c) Expression of Oprm1 in both acute and repeated NAc. (d) Expression of Fgf12 in both acute and repeated NAc. (e) Expression of Oprm1 in primary striatal culture. (f) Expression of Fgf12 in primary striatal culture.

**Supplementary Figure 6:**
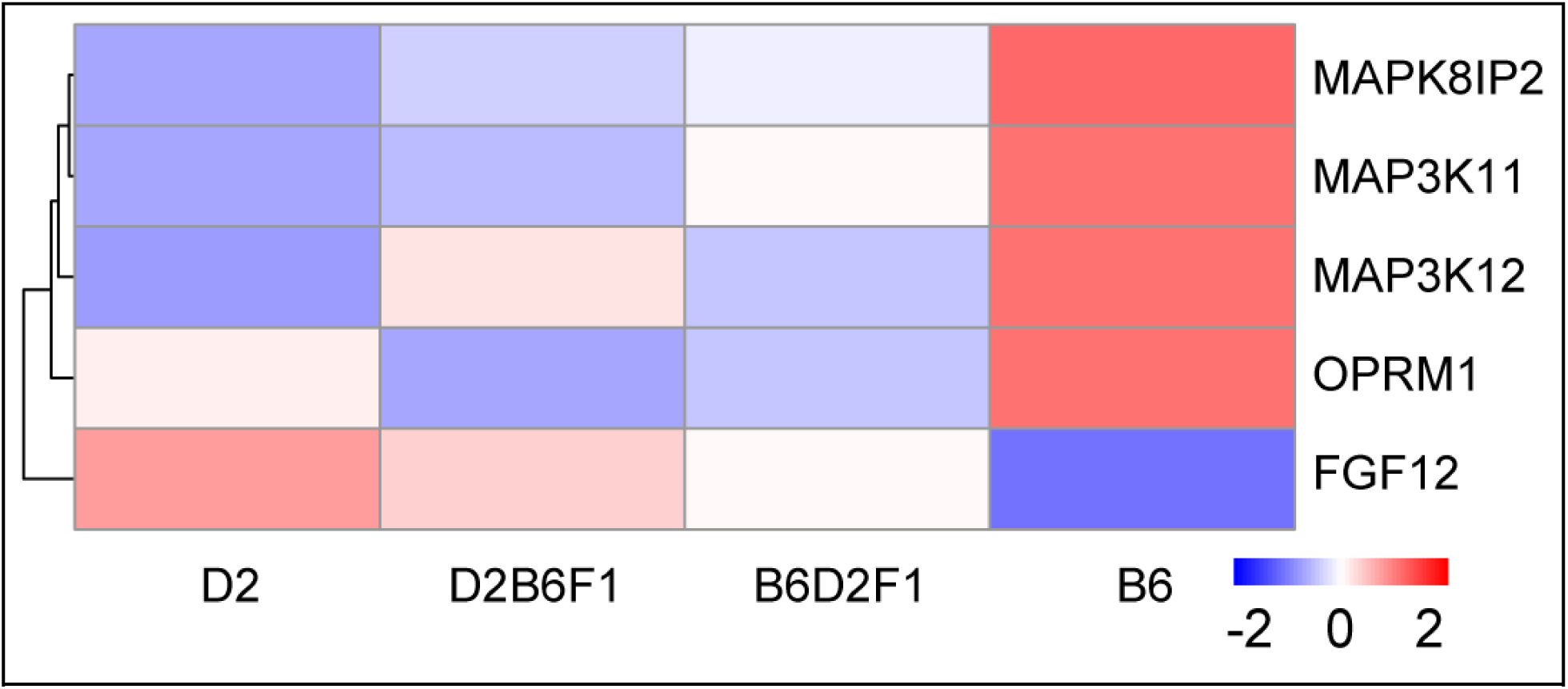
Levels of Oprm1, Fgf12, and MAP kinases proteins in the brains of BXD parental strains and F1s. Effect of genotype on associated MAP kinases and genes for both parental strains and F1 generations. The genotype effect shows the B allele is associated with high expression of Oprm1 and the MAP kinases Mapk8ip2, Map3k11, and Map3k12, but low expression of Fgf12, when compared to the D allele.

**Supplementary Figure 7:**
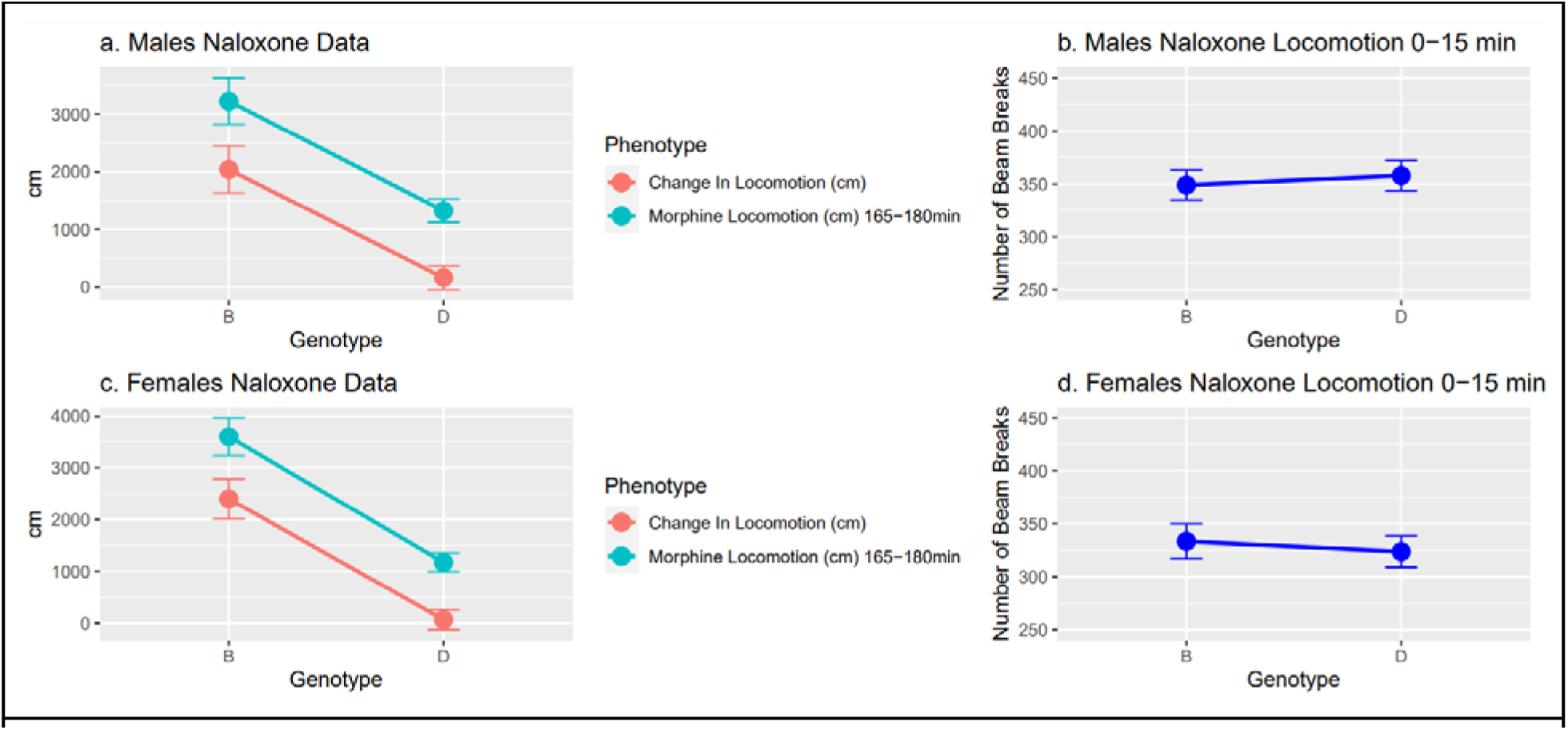
Naloxone Phenotypes vs. Genotypes. The turquoise points show morphine-induced locomotion (distance traveled in cm) between 165–180 min after injection. Orange points show the change in locomotion measured by the last 15 min of morphine locomotion response minus the first 15 min after naloxone injection. (a) Effect of genotype on two naloxone phenotypes in males. (b) Effect of genotype on number of beam breaks in an open field 0–15 min after naloxone injection in males. (c) Effect of genotype on two naloxone phenotypes in females. (d) Effect of genotype on the number of beam breaks in an open field 0–15 min after naloxone injection in females.

---

**Supplementary Table 1.** Genome-wide significant loci associated with morphine and naloxone responses (quantile-normalized)

| Locus | Peak LOD | Chr | Peak | Add Effect | Short description | Time | Sex ‡ | N Strains | GN BXD Trait |
| --- | --- | --- | --- | --- | --- | --- | --- | --- | --- |
| *Mor1a* | 3.99 | 1 | 74.07 | -0.40 | Morphine-induced locomotion, distance traveled | 0-15 | F | 64 | 11583 |
| *Mor1a* | 3.95 | 1 | 74.07 | -0.37 | Morphine-induced locomotion, distance traveled | 15-30 | F | 64 | 11586 |
| *Mor1a* | 4.42 | 1 | 77.64 | -0.42 | Morphine-induced locomotion, distance traveled | 0-15 | F | 64 | 11583 |
| *Nalx1a* | 3.96 | 1 | 128.54 | -0.43 | Naloxone-induced withdrawal, salivation | NA | MF | 64 | 11875 |
| *Nalx1b* | 4.14 | 1 | 167.17 | -0.44 | Naloxone-induced withdrawal, salivation | NA | MF | 64 | 11875 |
| *Mor1b* | 4.32 | 1 | 173.53 | -0.38 | Morphine-induced locomotion, distance traveled | 150-165 | F | 64 | 11584 |
| *Mor4a* | 4.06 | 4 | 21.69 | -0.39 | Morphine-induced locomotion, distance traveled | 45-60 | M | 63 | 11331 |
| *Mor5a* | 3.84 | 5 | 9.85 | 0.41 | Morphine-induced locomotion, distance traveled | 60-75 | F | 64 | 11589 |
| *Mor5a* | 4.62 | 5 | 9.85 | 0.47 | Morphine-induced locomotion, distance traveled | 75-90 | F | 64 | 11590 |
| *Mor5a* | 4.49 | 5 | 9.85 | 0.48 | Morphine-induced locomotion, distance traveled | 90-105 | F | 64 | 11580 |
| *Mor5a* | 5.01 | 5 | 9.85 | 0.52 | Morphine-induced locomotion, distance traveled | 105-120 | F | 64 | 11581 |
| *Mor5a* | 4.11 | 5 | 9.85 | 0.47 | Morphine-induced locomotion, distance traveled | 120-135 | F | 64 | 11582 |
| *Mor5a* | 4.00 | 5 | 13.91 | 0.41 | Morphine-induced locomotion, distance traveled | 60-75 | F | 64 | 11589 |
| *Mor5a* | 3.80 | 5 | 13.91 | 0.40 | Morphine-induced locomotion, distance traveled | 45-60 | F | 64 | 11588 |
| *Mor5b* | 3.86 | 5 | 82.22 | 0.41 | Morphine-induced locomotion, distance traveled | 30-45 | F | 64 | 11587 |
| *Nalx6a* | 4.69 | 6 | 116.94 | -0.42 | Naloxone-induced withdrawal, horizontal activity | 0-15 | MF | 64 | 11871 |
| *Nalx6a* | 4.65 | 6 | 116.94 | -0.42 | Naloxone-induced withdrawal, locomotion | 0-15 | MF | 64 | 11870 |
| *Nalx6a* | 3.98 | 6 | 116.94 | -0.39 | Naloxone-induced withdrawal, horizontal activity | 0-15 | F | 64 | 11614 |
| *Nalx6a* | 4.39 | 6 | 116.94 | -0.41 | Naloxone-induced withdrawal, locomotion | 0-15 | F | 64 | 11613 |
| *Mor7a* | 3.88 | 7 | 84.15 | -0.46 | Morphine-induced locomotion, distance traveled | 90-105 | M | 63 | 11323 |
| *Mor7a* | 5.28 | 7 | 96.83 | -0.44 | Morphine-induced locomotion, distance traveled | 0-15 | M | 63 | 11326 |
| *Mor7a* | 3.89 | 7 | 96.83 | -0.39 | Morphine-induced locomotion, distance traveled | 30-45 | M | 63 | 11330 |
| **Mor7a** | 3.88 | 7 | 96.83 | -0.39 | Morphine-induced locomotion, distance traveled | 45-60 | M | 63 | 11331 |
| **Mor7a** | 4.15 | 7 | 104.15 | -0.48 | Morphine-induced locomotion, distance traveled | 45-60 | M | 63 | 11331 |
| **Mor7a** | 3.96 | 7 | 107.29 | -0.43 | Morphine-induced locomotion, distance traveled | 45-60 | M | 63 | 11331 |
| **Mor7a** | 3.95 | 7 | 114.58 | -0.34 | Morphine-induced locomotion, distance traveled | 30-45 | F | 64 | 11587 |
| **Mor9a** | 3.90 | 9 | 83.86 | 0.45 | Morphine-induced locomotion, distance traveled | 120-135 | F | 64 | 11582 |
| **Mor10a** | 9.46 | 10 | 4.94 | -0.58 | Morphine-induced locomotion, distance traveled | 60-75 | M | 63 | 11332 |
| **Mor10a** | 8.12 | 10 | 4.94 | -0.60 | Morphine-induced locomotion, distance traveled | 75-90 | M | 63 | 11333 |
| **Mor10a** | 4.02 | 10 | 4.94 | -0.45 | Morphine-induced locomotion, distance traveled | 120-135 | M | 63 | 11325 |
| **Mor10a/**<br>**Nalx10a** | 4.33 | 10 | 5.61 | -0.42 | Naloxone-induced withdrawal, change in locomotion | NA | M | 63 | 11339 |
| **Mor10a** | 10.53 | 10 | 5.64 | -0.59 | Morphine-induced locomotion, distance traveled | 30-45 | M | 63 | 11330 |
| **Mor10a** | 10.06 | 10 | 5.64 | -0.59 | Morphine-induced locomotion, distance traveled | 45-60 | M | 63 | 11331 |
| **Mor10a** | 4.23 | 10 | 8.90 | -0.40 | Morphine-induced locomotion, distance traveled | 0-15 | F | 64 | 11583 |
| **Mor10a** | 3.80 | 10 | 8.90 | -0.42 | Morphine-induced locomotion, distance traveled | 120-135 | F | 64 | 11582 |
| **Mor10a** | 4.07 | 10 | 8.90 | -0.41 | Morphine-induced locomotion, distance traveled | 0-15 | M | 63 | 11326 |
| **Mor10a** | 5.05 | 10 | 8.90 | -0.51 | Morphine-induced locomotion, distance traveled | 120-135 | M | 63 | 11325 |
| **Mor10a** | 4.60 | 10 | 8.90 | -0.46 | Morphine-induced locomotion, distance traveled | 105-120 | F | 64 | 11581 |
| **Mor10a** | 6.86 | 10 | 8.90 | -0.59 | Morphine-induced locomotion, distance traveled | 105-120 | M | 63 | 11324 |
| **Mor10a** | 8.52 | 10 | 8.90 | -0.54 | Morphine-induced locomotion, distance traveled | 30-45 | F | 64 | 11587 |
| **Mor10a** | 6.44 | 10 | 8.90 | -0.52 | Morphine-induced locomotion, distance traveled | 90-105 | F | 64 | 11580 |
| **Mor10a** | 9.30 | 10 | 8.90 | -0.56 | Morphine-induced locomotion, distance traveled | 15-30 | M | 63 | 11329 |
| **Mor10a** | 7.79 | 10 | 8.90 | -0.52 | Morphine-induced locomotion, distance traveled | 15-30 | F | 64 | 11586 |
| **Mor10a** | 9.60 | 10 | 8.90 | -0.59 | Morphine-induced locomotion, distance traveled | 45-60 | F | 64 | 11588 |
| **Mor10a** | 9.58 | 10 | 8.90 | -0.59 | Morphine-induced locomotion, distance traveled | 60-75 | F | 64 | 11589 |
| **Mor10a** | 8.34 | 10 | 8.90 | -0.58 | Morphine-induced locomotion, distance traveled | 75-90 | F | 64 | 11590 |
| **Mor10a** | 7.51 | 10 | 8.90 | -0.61 | Morphine-induced locomotion, distance traveled | 90-105 | M | 63 | 11323 |
| **Mor10a** | 3.93 | 10 | 12.85 | -0.43 | Morphine-induced locomotion, distance traveled | 90-105 | F | 64 | 11580 |
| **Nalx12a** | 3.83 | 12 | 27.59 | 0.39 | Naloxone-induced withdrawal, locomotion | 0-15 | F | 64 | 11613 |
| **Nalx12a** | 4.38 | 12 | 27.59 | 0.44 | Naloxone-induced withdrawal, locomotion | 0-15 | MF | 64 | 11870 |
| **Nalx12a** | 4.91 | 12 | 27.59 | 0.46 | Naloxone-induced withdrawal, horizontal activity | 0-15 | MF | 64 | 11871 |
| **Nalx12a** | 4.26 | 12 | 36.01 | 0.47 | Naloxone-induced withdrawal, horizontal activity | 0-15 | F | 64 | 11614 |
| **Mor12a** | 4.03 | 12 | 70.75 | -0.42 | Morphine-induced locomotion, distance traveled | 0-15 | F | 64 | 11583 |
| **Mor12b** | 5.15 | 12 | 99.58 | -0.45 | Morphine-induced locomotion, distance traveled | 0-15 | F | 64 | 11583 |
| **Mor12b** | 5.44 | 12 | 99.58 | -0.43 | Morphine-induced locomotion, distance traveled | 15-30 | F | 64 | 11586 |
| **Mor12b** | 5.71 | 12 | 99.58 | -0.44 | Morphine-induced locomotion, distance traveled | 30-45 | F | 64 | 11587 |
| **Mor12b** | 5.60 | 12 | 99.58 | -0.43 | Morphine-induced locomotion, distance traveled | 65-70 | F | 64 | 11589 |
| **Mor12b** | 3.94 | 12 | 99.58 | -0.39 | Morphine-induced locomotion, distance traveled | 75-90 | F | 64 | 11590 |
| **Mor12b** | 4.01 | 12 | 104.28 | -0.40 | Morphine-induced locomotion, distance traveled | 15-30 | F | 64 | 11586 |
| **Nalx14a** | 4.00 | 14 | 52.13 | -0.41 | Naloxone-induced withdrawal, number of jumps | NA | M | 63 | 11336 |
| **Nalx14b** | 4.25 | 14 | 118.46 | -0.24 | Naloxone-induced withdrawal locomotion | 0-15 | MF | 64 | 11870 |
| **Nalx14b** | 4.36 | 14 | 118.46 | -0.25 | Naloxone-induced withdrawal, horizontal activity | 0-15 | MF | 64 | 11871 |
| **Mor16a/**<br>**Nalx16a** | 5.34 | 16 | 27.45 | -0.47 | Naloxone-induced withdrawal, change in locomotion | NA | M | 63 | 11339 |
| **Mor16a/**<br>**Nalx16a** | 10.56 | 16 | 27.45 | -0.61 | Morphine-induced locomotion, distance traveled | 165-180 | F | 64 | 11585 |
| **Mor16a/**<br>**Nalx16a** | 4.28 | 16 | 27.45 | -0.45 | Morphine-induced locomotion, distance traveled | 165-180 | M | 63 | 11328 |
| **Mor16a/**<br>**Nalx16a** | 5.83 | 16 | 28.99 | -0.51 | Morphine-induced locomotion, distance traveled | 150-165 | F | 64 | 11584 |
| **Mor16a/**<br>**Nalx16a** | 3.92 | 16 | 28.99 | -0.44 | Morphine-induced locomotion, distance traveled | 165-180 | M | 63 | 11328 |
| **Nalx16b** | 4.02 | 16 | 59.72 | 0.42 | Naloxone-induced withdrawal, salivation | NA | MF | 64 | 11875 |

‡ MF = male and female joint mapping. M / F = significant in both sexes but –logP value applies to first entry

**Supplementary Table 2.** List of Locomotion Trait IDs.

| Sex | Time (min) | TraitID |
| --- | --- | --- |
| females | 0-15 | BXD_11583 |
| females | 15-30 | BXD_11586 |
| females | 30-45 | BXD_11587 |
| females | 45-60 | BXD_11588 |
| females | 60-75 | BXD_11589 |
| females | 75-90 | BXD_11590 |
| females | 90-105 | BXD_11580 |
| females | 105-120 | BXD_11581 |
| females | 120-135 | BXD_11582 |
| females | 150-165 | BXD_11584 |
| females | 165-180 | BXD_11585 |
| males | 0-15 | BXD_11326 |
| males | 15-30 | BXD_11329 |
| males | 30-45 | BXD_11330 |
| males | 45-60 | BXD_11331 |
| males | 60-75 | BXD_11332 |
| males | 75-90 | BXD_11333 |
| males | 90-105 | BXD_11323 |
| males | 105-120 | BXD_11324 |
| males | 120-135 | BXD_11325 |
| males | 150-165 | BXD_11327 |
| males | 165-180 | BXD_11328 |

